# Axial segmentation by iterative mechanical signaling

**DOI:** 10.1101/2023.03.27.534101

**Authors:** Susan Wopat, Priyom Adhyapok, Bijoy Daga, Janice M. Crawford, Brianna Peskin, James Norman, Jennifer Bagwell, Stephanie M. Fogerson, Stefano Di Talia, Daniel P. Kiehart, Patrick Charbonneau, Michel Bagnat

## Abstract

In bony fishes, formation of the vertebral column, or spine, is guided by a metameric blueprint established in the epithelial sheath of the notochord. Generation of the notochord template begins days after somitogenesis and even occurs in the absence of somite segmentation. However, patterning defects in the somites lead to imprecise notochord segmentation, suggesting these processes are linked. Here, we reveal that spatial coordination between the notochord and the axial musculature is necessary to ensure segmentation of the zebrafish spine both in time and space. We find that the connective tissues that anchor the axial skeletal musculature, known as the myosepta in zebrafish, transmit spatial patterning cues necessary to initiate notochord segment formation, a critical pre-patterning step in spine morphogenesis. When an irregular pattern of muscle segments and myosepta interact with the notochord sheath, segments form non-sequentially, initiate at atypical locations, and eventually display altered morphology later in development. We determine that locations of myoseptum-notochord connections are hubs for mechanical signal transmission, which are characterized by localized sites of deformation of the extracellular matrix (ECM) layer encasing the notochord. The notochord sheath responds to the external mechanical changes by locally augmenting focal adhesion machinery to define the initiation site for segmentation. Using a coarse-grained mathematical model that captures the spatial patterns of myoseptum-notochord interactions, we find that a fixed-length scale of external cues is critical for driving sequential segment patterning in the notochord. Together, this work identifies a robust segmentation mechanism that hinges upon mechanical coupling of adjacent tissues to control patterning dynamics.

## Introduction

In vertebrates, the skeletal muscles, connective tissues, and bony elements (vertebrae and arches) of the axial skeletal system exhibit a segmented pattern that is established from the anteroposterior (AP) division of the embryo into a series of functional units called somites (Pourquié, 2011). Somite segmentation is regulated through a highly conserved genetic network of oscillating signals, referred to as the segmentation clock (Pourquié, 2003). When the clock genes are disrupted, a wide range of axial musculoskeletal defects are observed (Eckalbar et al., 2012; Sparrow et al., 2010; Whittock et al., 2004). In amniote embryos (mammals, birds and reptiles), perturbations to the segmentation clock impact patterning of the skeletal muscles, tendons, and vertebral column, particularly vertebral body and arch morphology (Aoyama & Asamoto, 1988; Stern & Keynes, 1987). By contrast, in teleost fish, clock mutations cause defects in muscle and myoseptum segmentation as well as vertebral arch morphology, whereas vertebral body malformations are comparatively mild (Fleming et al., 2004; Lleras Forero et al., 2018; van Eeden et al., 1998). Underlying these phenotypic differences between vertebrates are distinct developmental mechanisms that emerged in teleost fish and amniotes. Unlike amniotes, which predominantly rely on the re-segmentation of somite-derived sclerotome cells to pattern vertebral bodies, teleosts (modern bony fish) form vertebrae in a notochord-dependent manner via the production of metameric mineralized rings, known as chordacentra (Fleming et al., 2004; Gadow & Abbott, 1895). Chordacentra then serve as template for osteoblast recruitment and vertebral body development (Wopat et al., 2018). When the segmented template of the zebrafish notochord is missing, osteoblasts are still recruited to the notochord via the somite boundaries, producing hemi-vertebrae and diplospondyly (i.e., two vertebral elements per segment) (Peskin et al., 2020). This phenotype resembling the vertebral pattern of stem teleosts and primitive sarcopterygian lineages, the ancestors of teleosts and tetrapods, suggests that the morphogenetic program of the fish spine retains a core segmental program that is common to that of amniotes.

During vertebral body development in zebrafish, regions of chordacentra formation are first defined by segmentation of the notochord sheath, which patterns into cartilage-like and mineralizing domains in a sequential manner along the AP axis (Lleras Forero et al., 2018; Pogoda et al., 2018; Wopat et al., 2018). Notochord segments that underlie chordacentra express mineralizing proteins such as Entpd5a and Cyp26b1, while proteins characteristic of cartilage formation, including collagen proteins Col9a2 and Col2a1, define the domains that eventually develop into the intervertebral discs (IVDs). Sheath cells transition from a uniform *col9a2+* population to form alternating *entpd5a+* domains in a sequential fashion, with one segment forming every ∼8 hours in an A-to-P direction through continuously active Notch signaling (Wopat et al., 2018). This is in contrast to somitogenesis, which displays cyclic expression of Notch target genes (Liao et al., 2016; Özbudak & Lewis, 2008). Furthermore, while segment formation during somitogenesis is strictly linear (i.e., always sequential in the A-to-P direction), gaps in the segmented pattern of the notochord can be repaired with some delay (Lleras Forero et al., 2018; Wopat et al., 2018). Interestingly, previous work has also shown that mutations affecting somite boundary (SB) formation such as *fused somites* (*tbx6)* or a localized disruption of SBs during somitogenesis, impact the fidelity of notochord segmentation (Lleras Forero et al., 2018; Wopat et al., 2018). For example, experiments that transiently expose embryos to the Notch inhibitor DAPT during a short window of somite segmentation produce a gap in the pattern of SBs (Özbudak & Lewis, 2008). These DAPT gaps also lead to localized regions of notochord segmentation defects that coincide with the affected SBs (Wopat et al., 2018). In addition, transplantation experiments suggest that notochord-paraxial mesoderm interactions may also play a role in notochord patterning in amniotes (Ward et al., 2018). Together, these experiments strongly suggested that crosstalk between the notochord and the surrounding tissues has a role in regulating notochord segmentation in teleosts.

Here, we investigate the link between the spatial and temporal disorder that is observed during notochord segmentation in somite patterning mutants. We show that the interaction between the notochord sheath and the myosepta present at SBs leads to focal adhesion clustering ahead of notochord segment initiation with the activation of Notch and *entpd5a* at those sites. We find that when this spatial information is disrupted by genetic defects in SB formation or experimental severing of myosepta, notochord segmentation proceeds non-sequentially, indicating that locally provided mechanical cues at notochord-myoseptum contacts promote precise and sequential segmentation of the notochord. We captured these dynamics in a theoretical model which suggests that a fixed length scale of external cues is critical for driving sequential segment formation in the zebrafish notochord. This work uncovers a role for tissue-tissue interactions in sequential metameric patterning.

## Results

### Myoseptum-notochord interactions define the site of notochord segment initiation

Notochord segmentation still occurs in the absence of somite segmentation clock components (Lleras Forero et al., 2018) or when SBs are mispatterned (Wopat et al., 2018), albeit imprecisely. This loss of precision in the pattern of notochord segments in somite segmentation mutants and the correspondence of notochord segmentation defects to DAPT gaps suggested that SBs somehow influence notochord segmentation. To explore this connection, we first examined in detail the developing axial skeletal system (Figure 1A) using transgenic lines that specifically label the notochord sheath, its surrounding ECM, and the myosepta. To label the notochord ECM, we generated a functional knock-in fusion line for Calymmin, *TgKI(cmn-tdTomato)* using a CRISPR-based method we recently described (Levic et al., 2021). Calymmin is a zebrafish-specific extracellular matrix protein similar to Elastin (Peskin et al., 2020). Calymmin-tdTomato localizes to the outermost layer of the notochord extracellular matrix (ECM) (Figure 1B), as previously shown using immune-EM (Cerdà et al., 2002), where it is arranged as circumferentially-oriented fibers (Figure 1C). Imaging of 5 days post-fertilization (dpf) *TgKI(cmn-tdTomato)* larvae expressing *Tg(Bactin:pxna-eGFP)*, a transgene that labels the myotendinous junctions of myosepta, (Goody et al., 2010; Subramanian & Schilling, 2014) revealed a close association of the two tissues at dorsal and ventral locations of the notochord (Figure 1C and 1D). The myosepta also appear to infiltrate the Calymmin ECM layer of the notochord at the ventral locations (Figure 1E).

**Figure 1:**
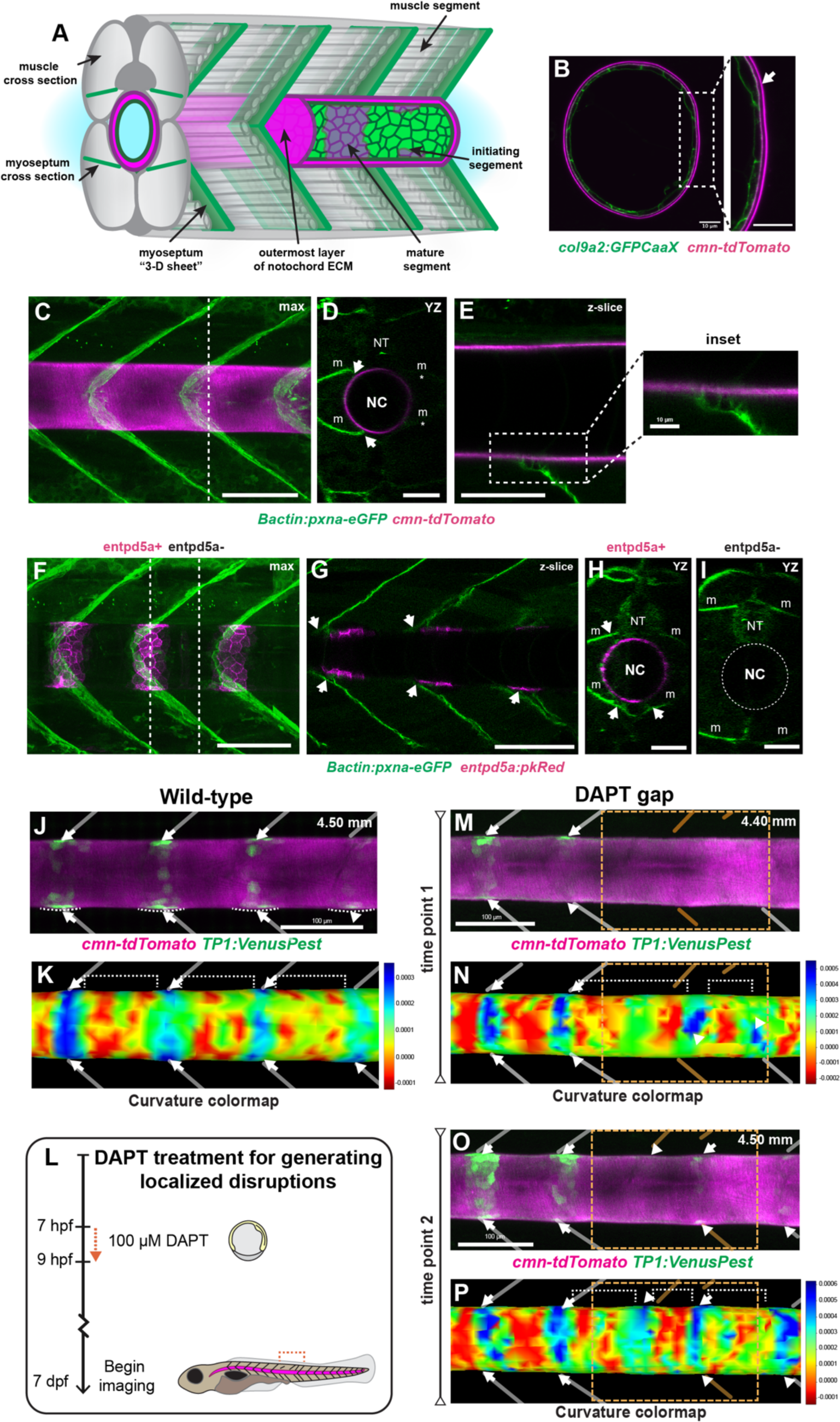
Myosepta contact and deform the outermost layer of the notochord sheath ECM in entpd5a+ domains. A) Schematic of developing axial musculoskeletal system in zebrafish. B) Cross-section of 5.0 mm standard length (SL) larvae expressing *TgKI(cmn-tdTomato)*, (magenta) which labels the outermost layer of the notochord ECM, and the notochord sheath reporter, *Tg(col9a2:GFPCaaX)* (green). Inset is zoomed in region of the cross section highlighting the collagen-rich gap between the outermost Calymmin-rich layer and the surface of the notochord. Scale bars are 10 µm. C) Max intensity projection of 4.50 mm SL larva expressing the notochord ECM reporter, *TgKI(cmn-tdTomato)* (magenta), and a reporter for the myosepta, *Tg(Bactin:pxna-eGFP)* (green). Dotted line corresponds to the optical YZ section depicted in (D). Scale bar = 100 µm. D) An optical YZ section of *TgKI(cmn-tdTomato)* and *Tg(Bactin:pxna-eGFP)* highlights the myosepta association with the outer ECM layer of the notochord (arrows) on the left side of the larva. Non-uniform illumination during image acquisition resulted in dimmer myosepta on the right side of the larva (denoted by *). Scale bar = 50 µm. E) A z-slice of a *TgKI(cmn-tdTomato)* and *Tg(Bactin:pxna-eGFP)* larva shows a ventral myoseptum infiltrates the ECM of the notochord. Scale bar = 50 µm. Inset is zooms in on connection. F) Max intensity projection of *Tg(Bactin:pxna-eGFP)* and notochord segment reporter *TgBAC(entpd5a:pkRed)* (magenta) in a 4.50 mm SL larva demonstrates that the myospeta (m) connections coincide with segment formation (arrows). Dotted lines correspond to optical sections in (H) and (I). NT denotes neural tube. Scale bar = 100 µm. G) Z-slice of the same larva demonstrates that myosepta contacts are closest to the notochord at dorsal and ventral locations. Scale bar = 100 µm. H) Optical YZ section of an *entpd5a+* region in the same larva expressing *Tg(Bactin:pxna-eGFP)* and *TgBAC(entpd5a:pkRed)*. Myoseptum-notochord connections are specified by arrows. Scale bar = 50 µm I) Optical YZ section of an *entpd5a)* region demonstrates that the angled myosepta do not contact the surface of the notochord. In (H) and (I) the myosepta and neural tube are labeled with (m) and (NT), respectively. Scale bar = 50 µm. J) Confocal image of 4.50 mm larva expressing *cmn-tdTomato* and *TP1:VenusPest*. Dotted lines highlight locations of curvature changes. Arrows point to *TP1*+ regions, while the arrowhead points to a region beginning to segment. Tracings of myosepta locations were done using DIC images (not shown). Scale bar 100 µm. K) Colormap of computed Gaussian curvature along the surface of the mesh corresponding to the surface extracted with *cmn-tdTomato* fluorescent expression. Arrows point to the same *TP1*+ regions which correlate to regions of positive curvature (blue). Dotted brackets highlight regular spacing between regions of positive curvature. L) Schematic of DAPT treatment from 7 hours post fertilization (hpf) to 9 hpf. After 9 hpf, embryos are then removed from DAPT exposure and allowed to develop normally. Time course imaging of early notochord segmentation starts at 7 dpf. M) Confocal image of a DAPT-treated larva expressing *cmn-tdTomato* and *TP1:VenusPest*. The dotted orange box and orange myosepta tracings denote the location of embryonic segmentation disruption. Arrows point to *TP1*+ regions, while the arrowhead points to a region displaying presumptive TP1 expression. Scale bar 100 µm. N) Colormap of corresponding ECM curvature shows segments with positive curvature (blue) correspond to *TP1*+ regions. Deformation is delayed in the disrupted region (orange dashed box) and also appears in irregular locations, highlighted by the varying widths of the white brackets and white arrowheads. O) Second timepoint of the same larva imaged in (M) shows that *TP1*+ segments are starting to back-fill in the disrupted region denoted by arrows and arrowheads in dotted orange box. P) Corresponding colormap shows that positive curvature is eventually is detected in the disrupted region (arrowheads and arrows located within the dotted orange box), indicating that weaker attachments are slower to deform the ECM surface.

To examine where specific locations of myoseptum connections occur in relation to notochord segment formation, we combined *TgBAC(entpd5a:pkRed)*, which specifically labels mineralizing segments (Wopat et al., 2018), with *Tg(Bactin:pxna-eGFP)*. Notably, myoseptum-notochord connections were located at sites of notochord segment formation (Figure 1F-I). To investigate the dynamics of these connections, we imaged *cmn-tdTomato* together with the Notch activity reporter *TP1:VenusPest* (Ninov et al., 2012). Interestingly, time course imaging revealed that the surface of the ECM deforms at locations of segment formation marked by the Notch activity reporter (Figure 1J). To quantitatively measure ECM deformation, we utilized the MATLAB toolbox, Image Surface Analysis Environment (ImSAnE) (Heemskerk & Streichan, 2015). This image processing approach identifies and extracts a surface of interest (SOI) from three-dimensional datasets, thus enabling us to examine the surface geometry of the notochord. Using our detected surface, we then mapped changes in curvature along the notochord sheath ECM (Figure 1K, Figure S1). In wild-type larvae, domains of active Notch signaling acquire a positive curvature, indicating these regions have bulged outward, while non-segmenting regions are mostly flat or display negative curvature values. Furthermore, changes in curvature appear to precede the activation of segmented Notch in sheath cells (Figure 1K).

To test if the ECM deformation depends on myosepta attachments, we generated regions of disrupted SBs flanked by normally segmented regions (DAPT gaps) using a previously established method (Özbudak & Lewis, 2008) (Figure 1L). Time course imaging of 7 dpf larvae expressing *cmn-tdTomato* and *TP1:VenuPest* revealed that Notch activation was delayed within the DAPT gaps (Figure 1M) and surface deformation occurred in irregular locations (Figure 1N). When we imaged the same region at a later stage, we observed Notch activation and surface deformation still lagged within the DAPT gap, but was detected in unaffected, posterior regions (Figure 1O and 1P). Together, these data show that notochord segment formation occurs at sites where myosepta contact and deform the notochord surface.

### Myosepta provide a local mechanical signal for notochord segmentation

The delay in notochord deformation and Notch activation observed within DAPT gaps suggested that the myoseptum-notochord connections establish a spatial pattern for segment initiation. To characterize the impact of defects in myoseptum attachments on notochord segmentation, we examined the dynamics of segmentation in *Bactin:pxna-eGFP* and *entpd5a:pkRed* expressing larvae with DAPT gaps. Based on the pattern of *pxna-GFP* expression, DAPT gaps ranged from 4 to 7 segments and were flanked by correctly patterned regions on the anterior and posterior sides, as previously reported (Özbudak & Lewis, 2008) (Figure 2A). Plotting the mean intensities of *Bactin:pxna-eGFP* and *entpd5a:pkRed* along the AP axis revealed that notochord segmentation defects specifically coincide with regions of disrupted myosepta (Figure 2B), further highlighting that disrupted notochord segmentation is impacted locally, as we previously observed (Wopat et al., 2018). Time course imaging of larvae with DAPT gaps revealed that notochord segment formation is delayed and imprecise in regions with disrupted myosepta, resulting in alterations to segment morphology (Figure 2C). Next, to assess how the spatiotemporal segmentation dynamics are impacted within and outside the disrupted areas, we shortened our image acquisition interval to 24 hours to enable tracking of individual *entpd5a+* segments in larvae from 5 to 12 dpf (Figure S2). We then quantified the patterning dynamics by assigning an ordering index to each segment. This index is a whole number which ranks each segment along the AP axis based on its initiation during segmentation (Materials and Methods, Figure 2D). To determine the ordering index for each larva, we compared the observed segmentation pattern against an expected sequential pattern. Additionally, to account for variations in DAPT gap sizes, we rescaled the segment positions so that the segments at the left and right boundaries of the disruption were labelled at −1 and 1 along the AP axis respectively. In both WT and undisrupted regions, we found that the ordering index showed little deviation from an expected sequential order (Figure 2E and 2F). However, in regions corresponding to disrupted myosepta, sequential segment initiation was lost and characterized by segments skipping ahead (positive mismatches) or filling in (negative mismatches) (Figure 2F). Collectively, these data indicate that the effect of gaps in myoseptum-notochord contacts are local.

**Figure 2:**
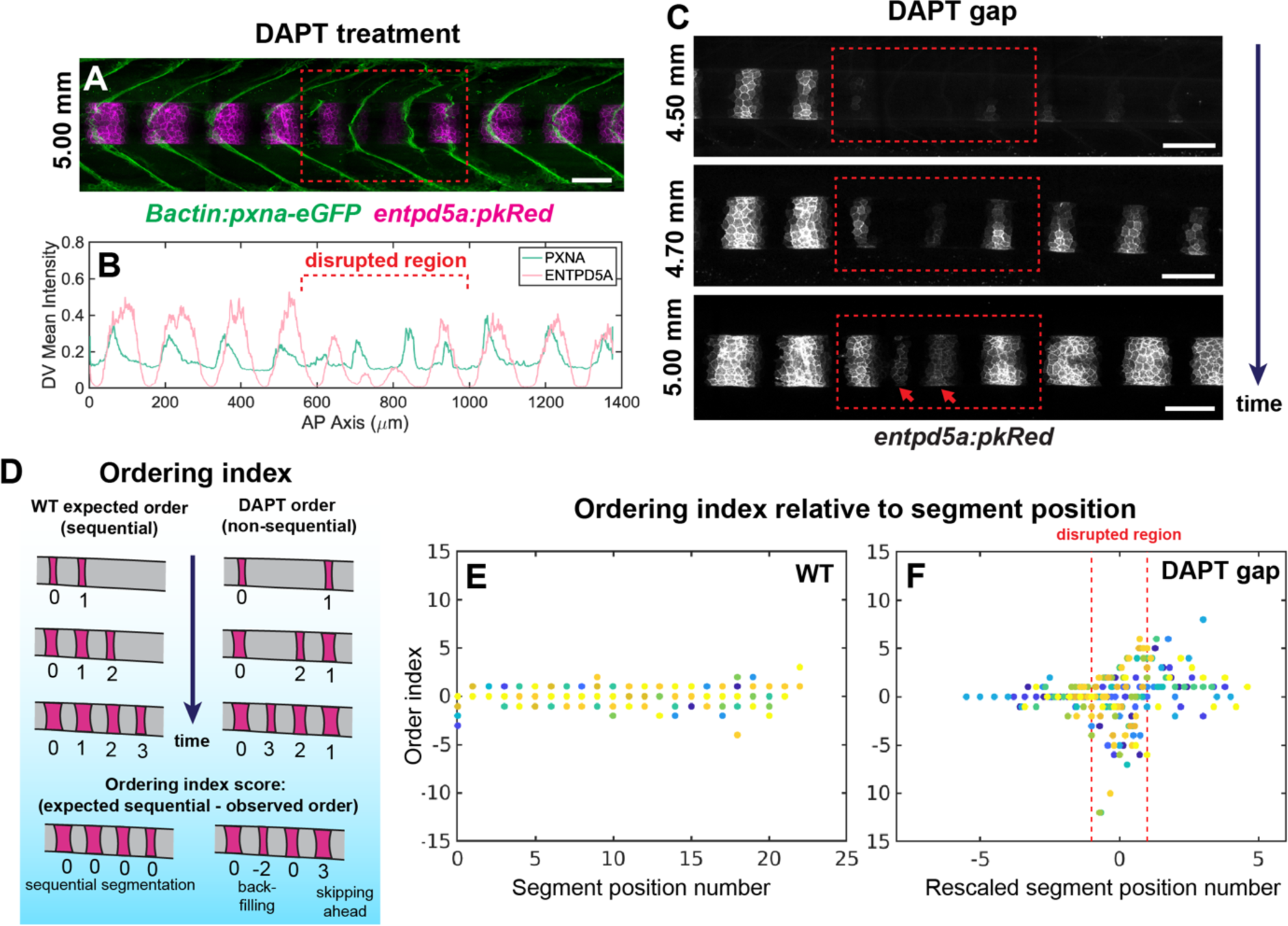
Disruption of somite segmentation correlates to defects in notochord patterning. A) Representative image of an DAPT-treated larvae expressing *Tg(Bactin:pxna-eGFP)* and *TgBAC(entpd5a:pkREd)*. Dashed red box signifies the location of somite boundary and subsequent myosepta disruption. Scale bar 100 µm. B) Trace plot depicting mean intensities for *Bactin:pxna-eGFP* and *entpd5a:pkRED* expression from image in panel (A). Well-defined peaks of *Bactin:pxna-eGFP* and *entpd5a:pkRED* fluorescent intensity are lost in the disrupted region. C) Time course of *entpd5a:pkRed* segmentation of the same larva shown in (A). Dashed red box highlights disrupted region and red arrows indicate segments back-filling in the skipped region. D) Schematic depicting criteria used to determine *entpd5a:pkRed* segment mismatch in wild-type and DAPT treated embryos. After ordering indexes are assigned to each segment, the final mismatch is calculated by subtracting the observed order from the expected sequential order. E) Plot depicting order mismatch values in along the AP axis of wild-type larvae. F) Plot of DAPT-treated larvae exhibiting regions both undisrupted and disrupted regions (denoted by dashed red lines). Due to variation in the size of the disrupted region from larva to larva, segment positions were used to align disrupted regions across individuals to allow for comparison. Individual larvae are represented as different color dots.

To directly test the role of myosepta in notochord segmentation, we adapted a laser ablation injury approach to disrupt myosepta without impacting the notochord (Kiehart et al., 2000, 2006). To disrupt a single segment, we laser ablated the four, connecting vertical myosepta (dorsal-left, dorsal-right, ventral-left, ventral-right) at 7 dpf (Figure 3A and 3D). Injuries to the myosepta were performed in focal planes well above the notochord in *Bactin:pxna-eGFP* and *entpd5a:pkRed* expressing larvae, posterior to the last detectable notochord segment, as determined by fluorescent microscopy. We confirmed the specificity of our laser injuries by using the vasculature reporter *flk:mCherry* and found that blood vessel morphology was unperturbed following cuts along the myosepta (Figure S3D-G). Immediately following laser ablation, we observed that cut myosepta recoiled and relaxed, indicating that tension along the myosepta was significantly reduced (Figure 3B and 3C). Laser ablated larvae were then allowed to recover for 2 days before performing an imaging time course (Figure 3E). At 2 days post laser cutting (dplc), *entpd5a+* expression was skipped in 15 out of 19 larvae in segments that directly neighbored the laser injured myoseptum (Figure 3E). Between 4 and 6 dplc, the injuries began to heal allowing formation of a notochord segment adjacent to the injured myoseptum to resume (Figure 3E).

**Figure 3:**
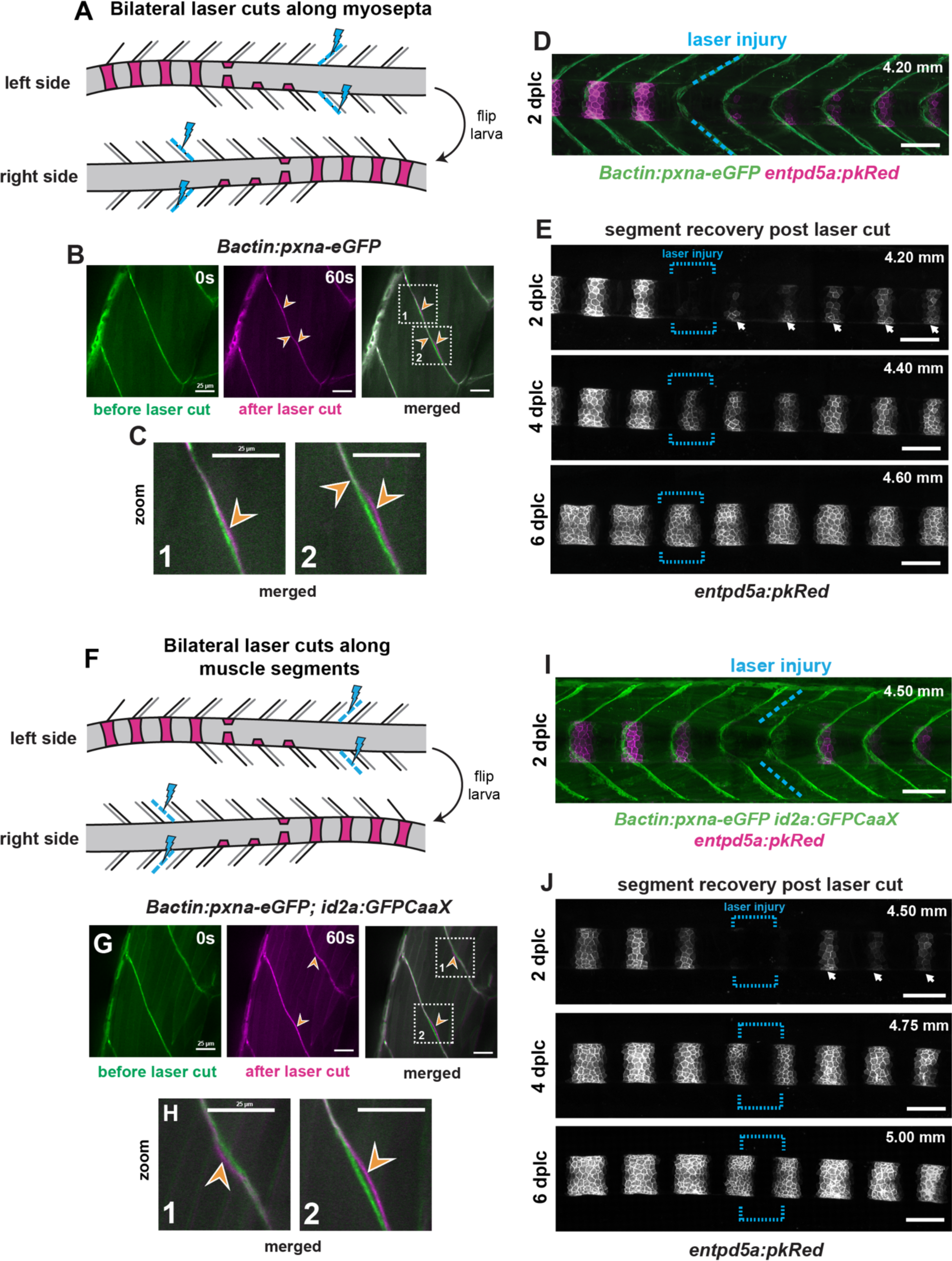
Laser injuries to the myosepta or axial muscles show segment initiation is regulated by force transmission at myoseptum-notochord connection points. A) Schematic illustrating the bilateral laser-cutting approach utilized to disrupt all four vertical myosepta (dorsal-left, dorsal-right, ventral-left, ventral-right). Injuries were always made in unsegmented regions in order to remove the segment initiation cue. B) Fluorescent images of focal plane used for laser-injury of *pxna-eGFP+* myoseptum in 7 dpf larva. The left-most image (green) is a representative image of the myoseptum pre-cut. The center image (magenta) is a representative image of the same myoseptum 60 s post-cutting. The right-most image (merged) is an overlay of pre-and post-cut images highlighting regions of “slackened” morphology following laser-injury (dotted boxes and orange arrowheads). Scale bar = 25 µm. C) Zoomed in perspective of dotted boxes from panel (B). Slackened morphology indicated by orange arrows. Scale bar = 25 µm. D) Max intensity projection of the same larva injured in panels (B) and (C) at 2 days post-laser-cutting (dplc). The dotted blue lines highlight the myosepta cuts. E) Time course of segment recovery post-injury at 2, 4, and 6 dplc. The region of laser-injury is indicated by dotted blue brackets. At 2 dplc, the adjacent notochord segment is skipped, whereas more posterior segments have already started forming (white arrows). At 4 dplc, the skipped segment from the injured area has recovered, but is delayed compared to more posterior segments. By 6 dplc, the skipped segment has fully recovered and is nearly indistinguishable from segments in neighboring, unaffected regions. Scale bars = 100 µm. F) Schematic illustrating the bilateral laser-cutting approach utilized to disrupt four adjacent muscle segments connected to two notochord segment regions via the bordering myosepta. G) Representative focal plane used for laser-cutting muscle segments in 7 dpf larva. Muscle cells are labeled with *id2a:GFPCaaX*, while the myosepta are still indicated by *Bactin:pxna-eGFP*. The left-most image (green) shows the muscle and myosepta morphology pre-cut. The center image (magenta) is a representative image of the same region 60 s post-cutting. The right-most image (merged) is an overlay of pre- and post-cut images, showing that both bordering myosepta are disrupted when the muscle is targeted (dotted boxes and orange arrowheads). Scale bars = 25 µm. H) Zoomed-in images of disrupted regions in the caudal (1) and rostral (2) bordering myosepta. Scale bars = 25 µm. Orange arrowheads indicate highly “slackened” regions. I) Max intensity projection of the same larva injured in panels (G) and (H) at 2 days post-laser-cutting (dplc). The dotted blue lines highlight muscle cuts. J) Time course of segment recovery following bilateral muscle cuts at 2, 4, and 6 dplc. The region of laser-injury is indicated by dotted blue brackets. At 2 dplc, two adjacent notochord segments were skipped, whereas more posterior segments have already started forming (white arrows). At 4 dplc, the skipped segments from the injured area have mostly recovered, but are delayed compared to more posterior segments. By 6 dplc, the skipped segment has fully recovered and is nearly indistinguishable from segments in neighboring, unaffected regions. Scale bars = 100 µm.

Next, to test whether forces transmitted through the myosepta are linked to notochord segment formation, we targeted muscle segments instead. To this end, we made laser cuts at the center of each surrounding muscle bundle (dorsal-left, dorsal-right, ventral-left, and ventral-right), posterior to the last detectable *entpd5a+* segment of the notochord (Figure 3F and 3I). Immediately following laser cutting, we observed that the two myosepta bordering the cut muscle segment displayed the same slackened morphology following direct targeting of a single myoseptum (Figure 3G and 3H). At 2 dplc, we observed that expression of *entpd5a* in the two notochord segments adjacent to the cut was skipped in 7 out of 8 larvae (Figure 3J). Having also observed myosepta penetrating at ventral attachment sites (Figure 1E), we specifically targeted these in a bilateral manner (ventral-left, ventral-right) (Figure S3A). Interestingly, when only ventral myosepta were cut, a delay in segment initiation was still observed, indicating that earliest segmentation signal is likely biased on the ventral side (Figure S3B and S3C). Together, the targeted laser injuries reveal that inputs from the axial musculature via myoseptum-notochord interactions locally instruct notochord segment initiation.

### Focal adhesions transmit mechanical signals from the myosepta

Our imaging and laser-cutting experiments suggested that segment initiation cues are mechanically transmitted to the notochord via the deformation of the ECM. We reasoned that cell-ECM complexes within the notochord sheath cells would be upregulated in response to these mechanical cues. To investigate this idea, we returned to our previously published RNA-sequencing dataset (Wopat et al., 2018) to identify candidate effectors of these interactions. We focused on the transitional or *entpd5a+/col9a2+* cell population, which corresponds to cells that initiate segment formation (Figure 4A). We found significant enrichment of transcripts for focal adhesion proteins including, *vcla*, *pxna*, *zyx*, *tln2a*, and *ptk2ab* (Figure 4B).

**Figure 4:**
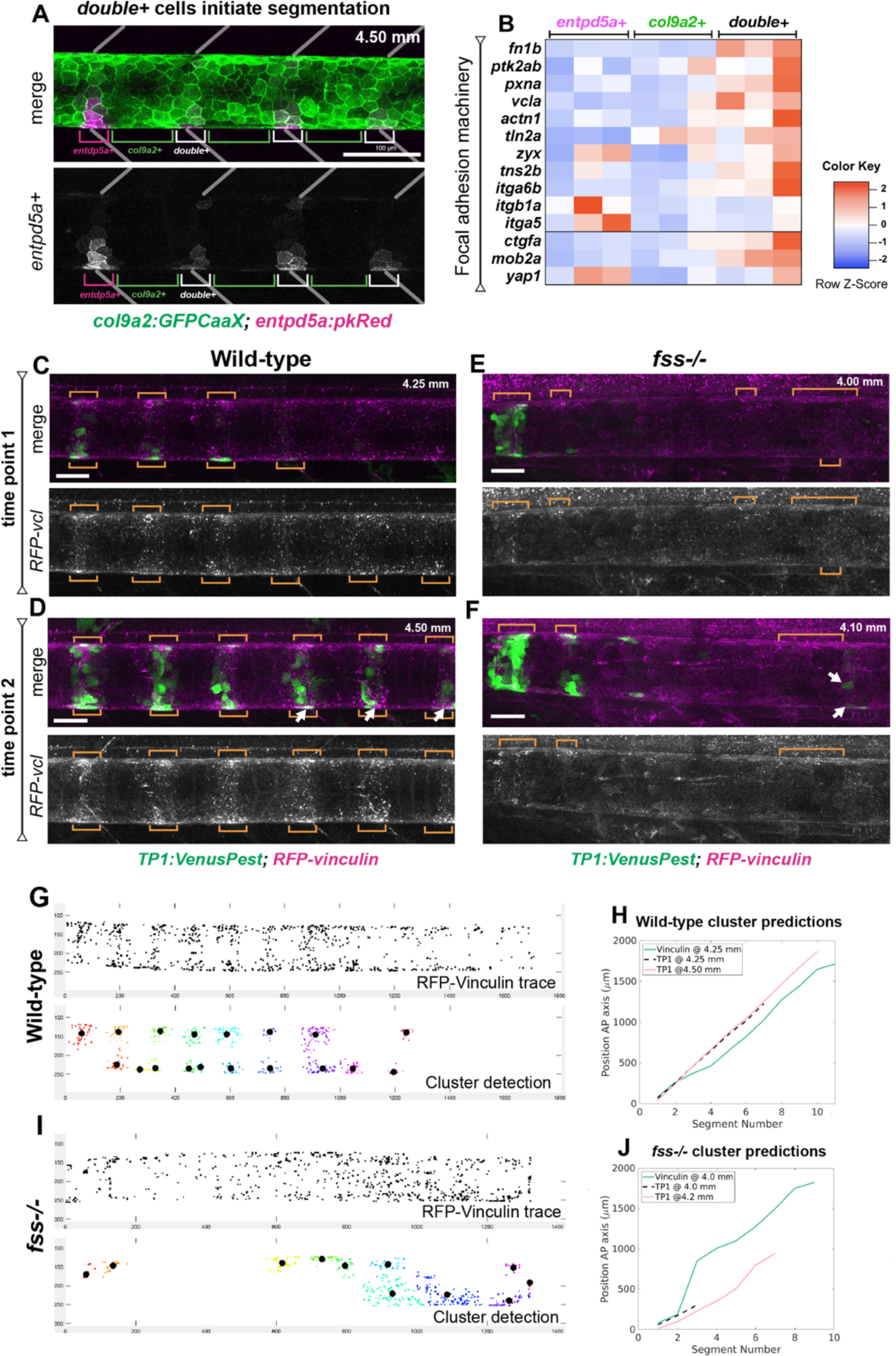
Myoseptum-notochord connections upregulate focal adhesion clustering to initiate transcriptional changes. A) Image of 4.50 mm SL larva expressing *col9a2:GFPCaaX* and *entpd5a:pkRed*. Magenta brackets highlight *entpd5a*+ populations, green brackets highlight *col9a2*+ populations, and white brackets point out *double*+ cells that were analyzed from previously RNA-sequencing analyses. Myosepta are outlined for reference. B) Heatmap of transcript enrichment profiles for *entpd5a+*, *col9a2+*, and *double+* populations. Candidates shown are known to be involved in focal adhesion complexes or have a known mechanosensing role. C) RFP-Vinculin localization in a 4.25 SL, *TP1:VenusPest* expressing, WT larva. Orange brackets point to dorsal and ventral clustering regions, which can be detected along the ventral side of the notochord ahead of *TP1:VenusPest* expression. D) Imaging of the same larva at a later developmental time point (4.50 SL) demonstrates that RFP-Vinculin puncta cluster together and correspond to newly formed *TP1:VenusPest* segments. E) Image of a *fss^-/-^* larva at 4.00 SL expressing *TP1:VenusPest* and injected with *RFP-vcla* mRNA. Orange brackets point toward regions that show some RFP-Vinculin puncta, but well-defined clusters are absent. F) A later time point of the same larva from (E) showing that TP1 expression is delayed and occurs in irregular locations (arrows). Few RFP-Vinculin puncta are present (denoted by orange brackets). Scale bars = 100 µm. G) Plot showing RFP-Vinculin puncta localization in the first WT time point. Below is a plot showing cluster detection (based on puncta number and proximity). H) Plot comparing Vinculin cluster localization from the first time point (4.25 mm SL) in blue, and *TP1* localization from the first and second time points (4.25 mm SL and 4.50 mm SL) in red and pink respectively. I) Plot showing RFP-Vinuclin puncta localization in the first *fss^-/-^*image. Below plot shows cluster detection (based on puncta number and proximity). J) Plot comparing Vinculin clusters to TP1 segment locations, indicating poor correlation between the earlier Vinculin clusters from time point 4.00 mm SL, to TP1+ segments at the second time point, 4.10 mm SL.

To determine where focal adhesions localize in sheath cells, we injected *RFP-vcla* cRNA into embryos expressing *TP1:VenusPest* at the one-cell stage and imaged larvae at 7 dpf (Figure 4C). Remarkably, RFP-Vinculin protein could be detected in the notochord sheath at 7 dpf and 2 days later, well beyond the typical life span of most cRNA translated proteins, indicating that Vinculin is highly stable in these cells (Figure 4C and 4D). RFP-Vinculin was present in puncta, likely corresponding to focal adhesions, and clustered at dorsal and ventral locations corresponding to myosepta attachment sites (Figure 4C). Interestingly, the cells displaying RFP-Vinculin clusters also expressed *TP1:VenusPest* at a later developmental timepoint, and displayed increased clustering of RFP-Vinculin as segments matured (Figure 4D).

Next, to assess if focal adhesion cluster assembly was dependent on myoseptum-notochord interactions, we used a mutation in *tbx6*-also known as *fused somites* (*fss*)-that fails to form well-defined embryonic somite boundaries and myosepta later in development (van Eeden et al., 1998). In *fss* mutants expressing *TP1:VenusPest*, RFP-Vinculin was only diffusely localized along the dorsal and ventral regions of the notochord (Figure 4E). At later timepoints, focal adhesion clustering was also diminished compared to WT (Figure 4F).

To determine how the observed focal adhesion clustering relates to notochord segment initiation, we developed a pipeline (Materials and Methods) to analyze the spatial and temporal dynamics of RFP-Vinculin clusters with respect to *TP1:VenusPest* segmentation. In WT larvae, periodic clusters of RFP-Vinculin were readily detected at the dorsal and ventral portions of the notochord, which corresponded to *TP1:VenusPest*+ cells at the subsequent developmental timepoint (Figure 4G). By contrast, RFP-Vinculin clusters in *fss* mutants were much more randomly localized and displayed decreased correlation with future *TP1:VenusPest*+ segments (Figure 4I and 4J). We also observed that some of the clusters were located laterally instead of being biased to the dorsal and ventral sides of the notochord (Figure 4I). These data indicate that myoseptum-notochord interactions lead to the formation of focal adhesions in sheath cells ahead of Notch activation and their differentiation into *entpd5a+* cells.

Focal adhesions have been shown to activate various mechano-transduction pathways that promote cell differentiation. To identify candidate effectors of this process we examined our RNA-sequencing data and found that both *yap1* and its target genes *ctgfa* and *mob2a* are enriched in the *entpd5a+/col9a2+* cell population (Figure 4B). To test the role of the Yap/Taz pathway in notochord segmentation, we used the QF2/QUAS system (Subedi et al., 2014) to express a dominant-negative form of Yap (DN-Yap) (Mateus et al., 2015) in *col9a2* cells. We found that overexpression of DN-Yap inhibited segmentation, as judged by the significant impairment in *TP1:VenusPest* induction (Figure S4). Together, these findings suggest that localized mechanical feedback at myoseptum-notochord connections on the dorsal/ventral portions of the notochord lead to focal adhesion assembly and the activation of the Notch-dependent segmentation program. Our results also suggest that Yap-mediated signaling plays a role in Notch activation in the notochord sheath.

### Disruptions in myosepta alter the spatial and temporal dynamics of notochord segmentation

Our analyses of focal adhesion clustering in WT and *fss* mutants reveal that myosepta define the positions at which *entpd5a* segments form. To better compare the full dynamics of notochord segmentation in wild-type and *fss* mutants, we took high resolution time courses of *entpd5a:pkRed* expression and computationally unwrapped 3-D images to visualize the entire surface of the notochord using ImSAnE (Heemskerk & Streichan, 2015) (Figure 5B and 5C, Figure S1). Surface projections revealed three phases of segmentation patterning in WT (Figure 5A). First, notochord segments begin as seeds located at regularly spaced intervals along the dorsal and ventral sides of the notochord. These seeds then grow and merge, forming a continuous ring, which later expands laterally to form mature segments (Figure 5B). By contrast, in *fss* mutants these seeds shift away from the dorsal and ventral portions of the notochord, often appearing on the lateral sides, and at irregular distances from adjacent seeds (Figure 5C). Computational extraction of these seed clusters and their positions around the notochord showed a shift in initiation sites from ventral locations in wild-type, to lateral ones in the *fss* mutants. (Materials and Methods, Figure 5D-F). These data are consistent with the position of focal adhesion clusters shown above (Figure 4). As segmentation proceeds, we observed that the early seeding defects compound to produce errors in segment merging and maturation (Figure 5B and 5C), with some seeds failing to merge and generating partial segments or segments of varying widths (Figure 5C).

**Figure 5:**
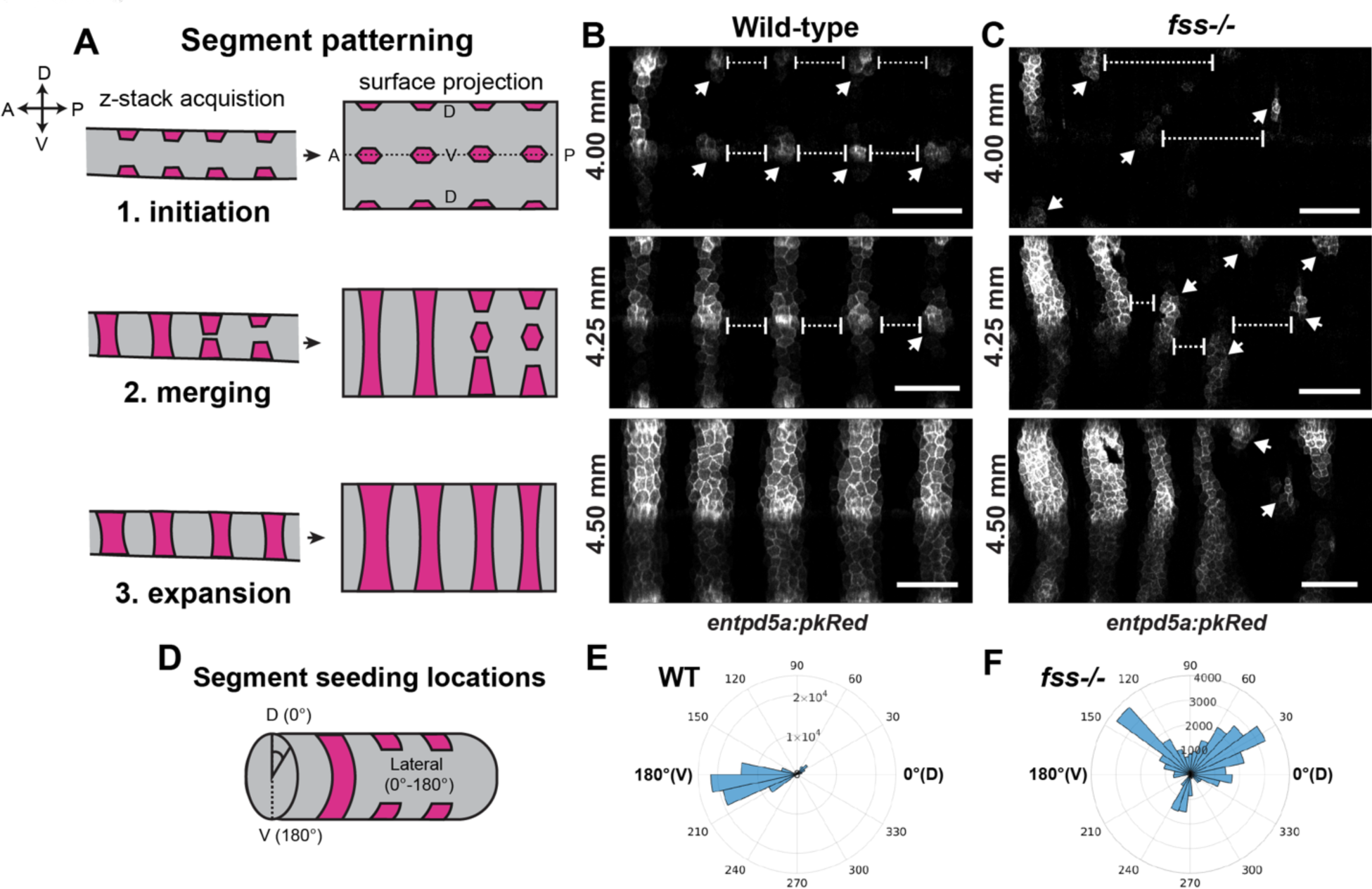
Myoseptum connections initiate segmentation at precise locations. A) Schematic highlighting the three major phases of notochord segmentation as visualized in a z-stack acquisitions and surface projections. B) In the first panel, surface projections of *entpd5a:pkRed* expression show wild-type segment initiation at 4.00 mm SL. Initiation is biased to the dorsal and ventral axes of the notochord (denoted by arrows). Spacing between segments is also tightly regulated (dotted brackets). At 4.25 dorsal and ventral seeds merge together to form a complete segment. Finally, at 4.50 segments begin to expand laterally. C) Surface projections of the *fss^-/-^* larva expressing *entpd5a:pkRed* show that at early stages (4.00 mm SL), segment seeds initiate along lateral portions of the notochord and at varying distances. During merging (4.25 mm SL), segment seeds fail to align with other seeds thereby generating partial segments or segments with altered morphology. These issues continue to compound at the expansion stage (4.50 mm SL) to produce segments of varying widths. Scale bars = 100 µm. D) Schematic showing seeding location analysis. E) Polar plot transforming the unwrapped positions around the notochord tube. Wild-type seeding locations are tightly controlled, typically occurring along the ventral portion of the notochord (180°). F) Seeding locations in *fss^-/-^* mutants show a shift toward lateral locations (30-150°), indicating that tightly regulated spatial control is lost without myoseptum instruction.

To investigate how spatial defects impact the temporal dynamics of notochord segmentation (Figure 6A and 6B), we imaged *entpd5a* expression in WT and *fss^-/-^* larvae every 24 hours for several days. Using our previously developed computational routine, we calculated ordering indexes and the mismatches relative to an ideal sequential pattern (Figure 2D). We found that deviations in segment formation mostly clustered around 0 for WT, whereas *fss* mutants lost sequential segmentation (Figure 6C). We also noted that on average it took WT larvae 11 days to fully segment, whereas *fss* mutants took 14 days, indicating that the patterning dynamics are significantly slowed when SBs/myosepta are disrupted. Using our ordering indexes and ranked sequence of segment formation, we also characterized the emergence of new segments over time by tracking inter-segment distance between two consecutively formed segments (Figure S5A). A value of 1 indicated segments were sequentially and proximately created. We found that WT values cluster around 1, whereas the *fss* mutants display a broad spread in the formation of subsequent segments (Figure 6D). Interestingly, monitoring the average distance between unranked segments in *fss^-/-^* mutants shows segments fill in to eventually achieve a periodic inter-segment distance comparable to that of WT (Figure 6E). These data show that the loss of regular spatial information from myosepta leads to slower, non-sequential segmentation dynamics within the notochord sheath. Our data also suggest the existence of a robust error correction process that fixes early patterning defects to produce a relatively well-segmented pattern even when the external input is highly disorganized.

**Figure 6:**
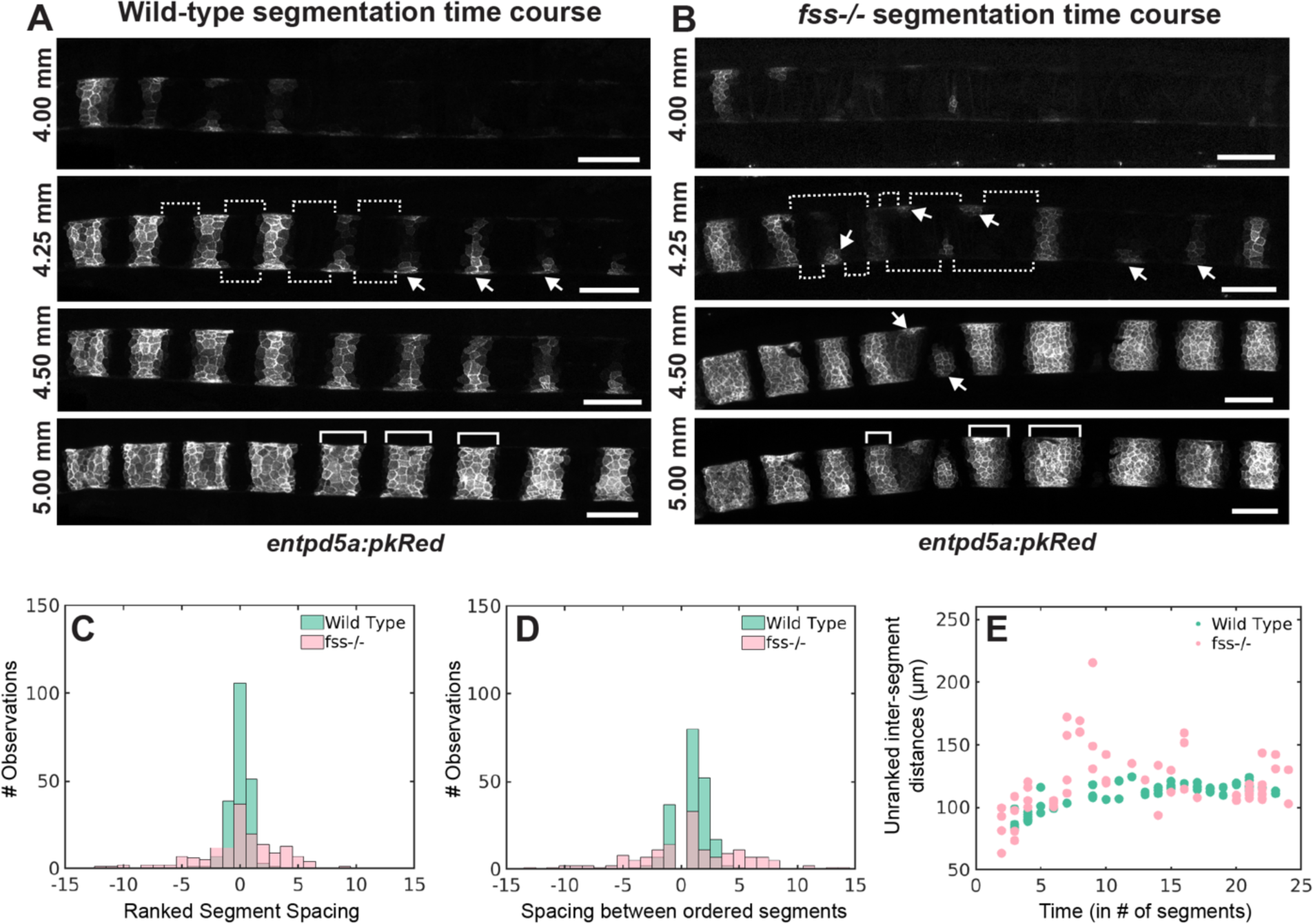
Notochord segmentation switches to an error-prone autonomous mechanism when instructive myoseptum signals are absent. A) Time course of a wild-type larva expressing *entpd5a:pkRed* imaged every 48 hours from 4.00mm to 5.00mm SL. White arrows point to segment initiation events, which occur posterior to fully formed segments. Dotted brackets represent distances between adjacent segments, and solid brackets show mature segment widths. B) Time course of a *fss^-/-^* larva expressing *entpd5a:pkRed* imaged every 48 hours from 4.00mm to 5.00mm SL. White arrows point to segment initiation events which often back-fill gaps. Dotted brackets represent distances between adjacent segments, and solid brackets show mature segment widths. Scale bars = 100 µm. C) Computed order mismatch observed in wild-type (n=9) and *fss* (n=8) larvae expressing *entpd5a:pkRed* (p value = 5.32e-07). D) Number of segments in between consecutively ranked segments in wild-type (n=9) and *fss* (n=8) larvae expressing *entpd5a:pkRed* (p value = 4.61e-07). E) Inter-segment distances observed in wild-type (n=9) and *fss* (n=8) larvae expressing *entpd5a:pkRed* over each time point without considering any ranking order. Distances between segments in the wild-type and *fss^-/-^* conditions converge once the notochord is close to being fully segmented.

### A theoretical model that captures the dynamics of notochord segmentation

Even when spatial cues are completely compromised as in *fss* mutants, the unsegmented notochord still manages to move toward a final segmented pattern (Figure 7A). However, the time evolution of this process shifts from a predominantly sequential mode to non-sequential when the spatial information is disorganized. We therefore investigated if the emergence of periodic stripes in the notochord and their assembly dynamics could be understood from minimal models which do not make any assumptions on the underlying molecular mechanisms a priori. Theoretically, several mechanisms including Turing’s reaction-diffusion system of morphogens (Gierer & Meinhardt, 1972; Turing, 1952) cell-cell interaction-based models (Nakamasu et al., 2009), and mechanical instabilities (Murray, 2003) could generate periodic patterns. All these mechanisms could also be understood within a uniform framework of model systems with competing short-range attraction and long-range repulsion (Hiscock & Megason, 2015a). The phase transition associated with such competing interactions gives rise to modulated phases in a variety of physical systems (Seul & Andelman, 1995), and in the context of biological systems these can be applied as models of non-equilibrium pattern formation (Gierer & Meinhardt, 1972; Kondo & Miura, 2010; Lavrentovich et al., 2016). The simplest model system obeying the requisite criterion can be captured by an effective free energy functional (Brazovskiǐ SA, 1975) with dynamics governed by the corresponding Swift-Hohenberg equation (Cross & Hohenberg, 1993; Swift & Hohenberg, 1977). We adapted these ideas to formulate a coarse-grained expression that describes the notochord system using a spatially anisotropic functional (F) for a scalar order parameter Φ that captures whether a cell has been activated by Notch (Φ>0) or not (Φ<0) (Supplementary Text). At sufficiently low noise (τ), this model favors the formation of spatially modulated striped domains of length scale λ=2π/*k*_0_, which represents the alternate domains expressing either *entpd5a* or *col9a2* along the AP (*x*) axis of the notochord. The effective model contains a term that favors interfaces perpendicular to the *x*-axis and one that inhibits domain formation perpendicular to the *y*-axis. The inherent anisotropy of the tubular form of the notochord makes such assumptions plausible. At low fluctuations, the functional is minimized by the creation of interfaces between cell domains of scale λ. The system dynamics show that for fixed *k*_0_, the system transitions from nucleation (i.e., random segment formation) to front propagation (i.e., sequential segment formation) as τ diminishes, with larger λ favoring front propagation (Figure 7B). Other mechanisms such as a parameter gradient in the coefficient of Φ^2^ term in the functional or adding production gradients in the dynamical equation could also orient stripes (Hiscock & Megason, 2015b) perpendicular to the AP axis. However, here we have not explored whether such processes would also reproduce the dynamics of the patterning process or if the underlying molecular mechanism could be potentially tested for those hypotheses.

**Figure 7:**
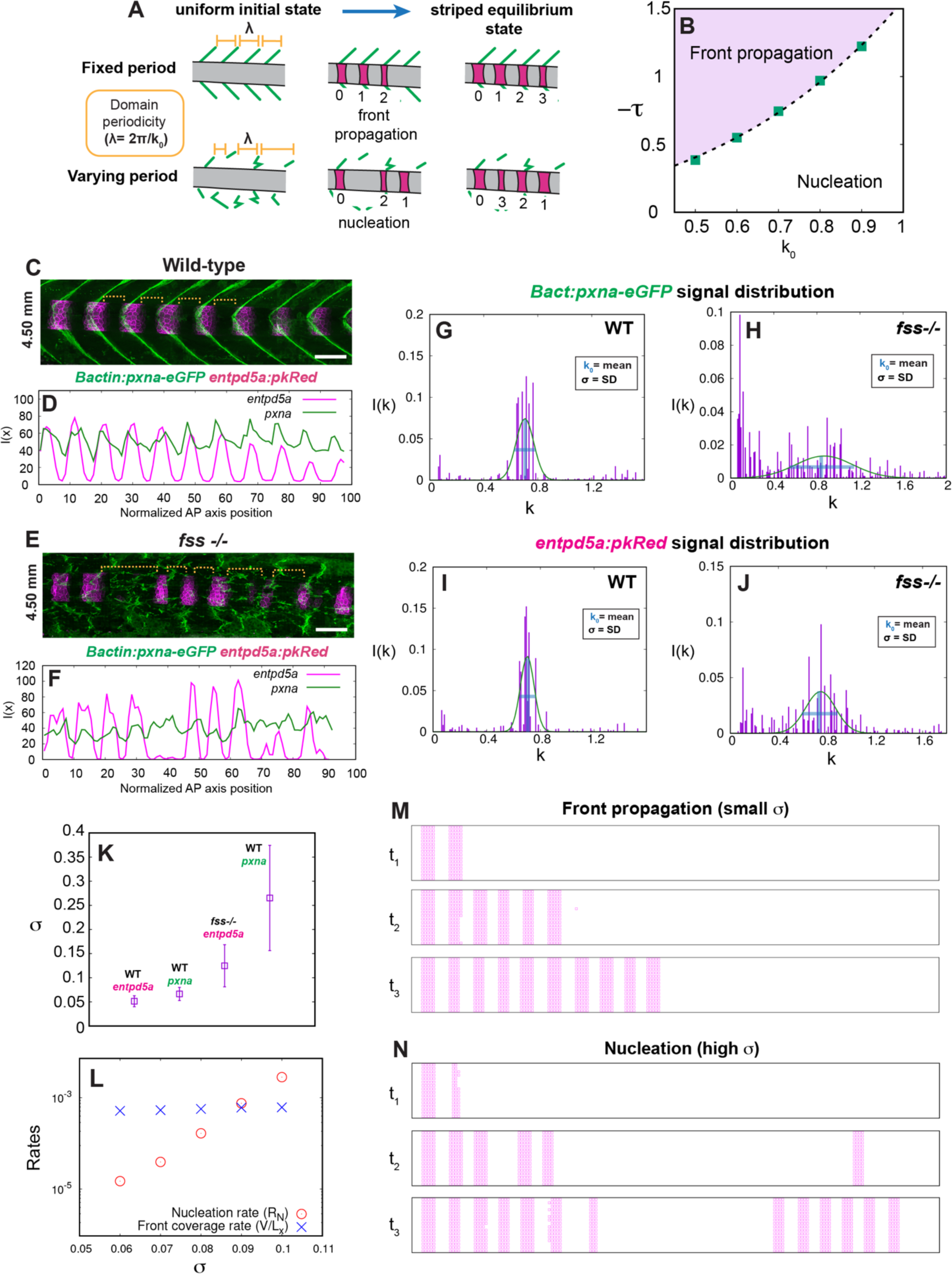
Spatial precision during notochord segmentation dictates temporal dynamics of segment initiation. A) Schematic of differing segmentation regimes when notochord-myoseptum connections are present at regular length-scales (fixed period) or noisy (varying period). B) Relaxation regimes in the k_0_-*τ* plane for *τ*<0 showing nucleation and front propagation as the two relaxation regimes of the system described by the free energy F_A_ to the striped phase starting from the initial seeded state. Here, L_x_=192, L_y_=16 and σ=0. C) Max projection of 4.50 mm SL wild-type larva expressing *Bactin:pxna-GFP*, and *entpd5a:pkRed*. Dotted brackets highlight regular spacing between adjacent segments. D) Trace plot depicting mean intensities for *Bactin:pxna-eGFP* and *entpd5a:pkRED* expression from WT image in panel (C). Well-defined signal peaks of *Bactin:pxna-eGFP* and *entpd5a:pkRED* fluorescent intensity occur at the same frequency. AP-axis is rescaled in units of mean cell width. E) Representative image of 4.50 mm SL *fss-/-* mutant expressing *Bactin:pxna-GFP*, and *entpd5a:pkRed*. Dotted brackets show variations in spacing between adjacent segments. F) Trace plot depicting mean intensities for *Bactin:pxna-eGFP* and *entpd5a:pkRED* expression from *fss^-/-^* image in panel (E). *Bactin:pxna-eGFP* fluorescent intensity is noisy along the AP axis, and does not exbibit signal peaks at a specific frequency. However, *entpd5a:pkRed* shows a segmented pattern, though its frequency is not regular. AP-axis is rescaled in units of mean cell width. G) Distribution of *Bactin:pxna-eGFP* signal from wild-type larva (n=6). H) Distribution of *Bactin:pxna-eGFP* signal from *fss*^-/-^ larva (n=6). I) Distribution of *entpd5a:pkRed* signal from wild-type larva (n=6). J) Distribution of *entpd5a:pkRed* signal from *fss*^-/-^ larva (n=6). K) Plot of the standard deviations (SDs) from the *entpd5a* and *pxna* FT signals in WT and *fss^-/-^* larvae. L) Numerical estimate of the front coverage rate (V/L_x_) and nucleation (R_N_) rates for different values of σ. For small σ, front propagation dominates, but nucleation takes over as σ increases. Here, *τ* = −0.80, k_0_ = 0.7. M) Typical temporal (t_1_<t_2_<t_3_) evolution of linearly spaced configurations in the front propagation regime with small values of σ. Domains form sequentially. (σ = 0.05, *τ* = −0.80, k_0_ = 0.7). N) Typical temporal (t_1_<t_2_<t_3_) evolution of linearly spaced configurations showing non-sequential ordering of domains in the nucleation regime for high values of σ. (σ = 0.12, *τ* = −0.80, k_0_ = 0.7).

To obtain experimental values for *k*_0_ we extracted fluorescent signals from WT and *fss^-/-^* larvae expressing *Bactin:pxna-GFP* and *entpd5a:pkRed* (Figure 7C-F). We then performed a Fourier transform (FT) of each signal and calculated the standard deviation (σ) and mean (*k_0_*) from the ensemble averaged distribution (n = 6 for WT and *fss^-/-^*) (Materials and Methods, Figure 7G-J). Plotting σ from the *entpd5a* and *pxna* FT signals from WT images, we found that their variance is relatively small and of similar range. By contrast, *fss^-/-^*larvae present a much higher variance for both markers (Figure 7K). Interestingly, we found that the *pxna* signal displays more variation compared to *entpd5a*, indicating that the pattern of myosepta is much more disorganized than the final segmented pattern of the notochord (Figure 7K).

We next used our model to analyze how the dynamics of the system changes with σ. By computing the nucleation rates and the front velocity (Supplementary Text), we found that for a given *τ*, the system undergoes a crossover from front propagation to nucleation as σ increases (Figure 7L). Simulations of the temporal dynamics of the system for the experimentally extracted values of σ and *τ* produce front propagation and nucleation regimes respectively (Figure 7M and 7N, Supp. Movies 1-3). For σ values obtained from WT, we observe that the dynamics is dominated by a sequential propagation (Supp. Movie 1); however, in some of the simulation runs, a small loss in order was also observed (Supp. Movie 2). Interestingly, similar defects are experimentally observed for WT fish in the rare occasions where a small loss of sequential order is detected (Figure S5B). In contrast, σ values derived from *fss* mutants show a much stronger loss in temporal order (Supp. Movie 3, Figure S5C). The experimental values for the temporal ordering of segments and domain width are compared with the simulation results and found to be in good agreement (Figure S5D-G). Intuitively, this behavior can be understood as the following: fluctuations in the parameter *k*_0_, lead to fluctuations in the energy barrier that must be overcome for nucleation, which in turn leads to disorder in patterning, as segments occasionally form at random locations.

Together, our phase-ordering model suggests that when inputs are provided at a fixed length scale, such as regularly spaced myosepta, sequential domain formation should be observed. However, fluctuations in the position of these inputs introduce variability in the timescale of nucleation and can result in periodic segments forming with spatial and temporal disorder. Consistent with this physical picture, the increase in spatial noise of external inputs found in the *fss* mutants delays patterning at notochord-myoseptum interaction sites with weak or diffuse signals, resulting in non-sequential segment initiation. Altogether, our study suggests that regularly spaced interactions with myosepta from the axial musculature contributes to precise and sequential segmentation in the notochord sheath.

## Discussion

In this study we report that mechanical coupling of the axial musculoskeletal tissues enables sequential and precise segmentation of the zebrafish notochord, thereby ensuring proper patterning of the vertebral column. While, previous work has demonstrated mechanical interactions between the notochord and paraxial mesoderm are critical for axis elongation (Guillon et al., 2020; McMillen & Holley, 2015; Tlili et al., 2019), we also find that axial tissue interactions provide an instructive role during notochord segmentation later in development. By the onset of notochord patterning, the embryonic somites have already differentiated into more specialized axial tissues such as muscle segments and tendon-like myosepta. The myoseptum contacts at the surface of the notochord ECM define the precise location where notochord segment seeds form, thereby linking early embryonic patterning to the later events of notochord segmentation. Furthermore, it seems possible that coordination between these patterning processes may provide a conserved mechanism for ensuring efficient musculoskeletal connectivity during development.

Using a combination of mutant, pharmacological, and laser ablation experimental approaches, we find that these myoseptum-notochord interactions provide a mechanical cue for segment initiation. Furthermore, there may be additional mechanisms that reinforce signaling at these attachment sites over time, such as progressive changes to the ECM at the interface of the myoseptum and notochord sheath (Grotmol et al., 2006). One element to consider is myoseptum maturation via the upregulation of Type-I Collagen (Wood & Currie, 2017), which coincides with the onset of notochord segmentation. An increase in rigidity at the dorsal and ventral myoseptum contact sites may locally impart mechanical cues from the growing muscle or maturing myosepta (Subramanian & Schilling, 2014; Tlili et al., 2019), to induce an increase in focal adhesion clustering and assembly. Mechanosensitive transcriptional changes, such as the upregulation of Yap signaling may then feed into the segmentation transcriptional program to initiate seeds. When the external signals from the myosepta are weak or irregular, as in *fss* mutants or DAPT-treated conditions, focal adhesions fail to cluster efficiently and notochord segmentation loses spatiotemporal precision.

Although the attachments between the myosepta and notochord evidently increase spatiotemporal precision of notochord segmentation, we find that even in WT larvae, segment initiation is not purely sequential. Our simulations revealed that temporal fluctuations and variance in the periodicity of myoseptum signals place the wild-type notochord close to the crossover point between sequential and non-sequential patterning (Figure 7B). This may provide the system with a robust mechanism that ensures segment formation even when part of the patterning information is lost. In *fss^-/-^* and DAPT-treated larvae, where spatial cues become especially noisy, the autonomous notochord segmentation program then becomes dominant. While this program is less accurate and produces defects that arise as early as the first initial seeding events, the system can eventually fill in skipped regions, or gaps lacking segments. This autonomous program also ensures that filled-in segments manage to move toward more uniform inter-segment spacing and produce similar inter-segmental distances as those of WT notochords (Figure 6E). Furthermore, the reduction in disorder between the pattern of myosepta (*pxna-GFP*) and *entpd5a* (Figure 7L) suggests that the notochord segmentation program is able to filter out some of the noise from poorly patterned myosepta to produce a relatively regular pattern. Together, these results suggest that an axial tissue-coupling mechanism dominates segment initiation, whereas autonomous notochord patterning mechanisms can drive segment growth and correct defects by back-filling when spatial signals are poorly defined.

Using a phase-ordering model, we suggest that when myoseptum cues are provided at discrete locations (single length-scale), spatial and temporal dynamics of segment initiation are preserved. By contrast, when external positional cues are irregular, temporal regulation of segment initiation is lost. Dissecting the molecular mechanisms that might give rise to the traveling wavefront and phase ordering dynamics predicted by the model will be a major goal of future research.

## Acknowledgements

We would like to thank Ken Poss for comments on the manuscript, Marcos Gonzalez-Gaitan and Olivier Pourquie for discussions, and the Duke Zebrafish Core Facility for fish care. We also thank Clarissa Henry for kindly providing the *Bactin:pxna-GFP* transgenic line and Sebastian Streichan for providing help with ImSAnE implementation. This work was funded by an HHMI Faculty Scholars Award to M.B., R01HL54737, an NIH MIRA R35GM127059 to D.P.K., and a DST grant from Duke University to S.D.

## Author contributions

S.W. and M.B., conceived and designed this project; S.W., P.A., J.C., B.P., J.N., J.B., and S.F. performed experiments and analyzed data; P.A. developed the original image analysis tools; B.D., S.D., D.P.K., and M.B. analyzed data. B.D., M.B., and P.C. conceived and generated the theoretical model. S.W. and M.B. wrote the manuscript with input from the other authors.

## Materials and Methods

### Fish stocks

Adult zebrafish of the Ekkwill strain (EK) were raised and bred as previously described (Ellis et al., 2013; Garcia et al., 2017) All experiments with animals were approved by Duke University. Individual fish were used for genetic manipulation experiments and compared to siblings and experimental control fish of similar size and age. Independent experiments were repeated using separate clutches of animals. Strains generated for this study: TgKI(cmn-tdTomato). Previously published strains: Tg(col9a2:QF2)pd1163, Tg(id2a:GFPCaaX)pd1167, TgBAC(entpd5a:pkRed)hu7478 (Wopat et al., 2018), Tg(TP1:VenusPEST)s940 (Ninov et al., 2012), Tg(Bactin:pxna-GFP) (Goody et al., 2010), Tg(flk:mCherry)s896 (Chi et al., 2008), and Tg(col9a2:GFPCaaX)pd1151 (Garcia et al., 2017). Transgenic lines were either generated using either the Tol2 system (Kawakami, 2007) or CRISPR knock-in strategies as previously described (Levic et al., 2021)

### Generating transgenic lines

**TgKI(cmn-tdtomato fusion line):** A PCR fragment containing 312bp of intronic sequence and exons 53/54 of *cmn* were inserted into a *pUC19-TgKI-MCS-tdTomato-polyA* vector backbone by cut and paste cloning. The following primers were used to amplify the C-terminal portion of *cmn*: BamHI_cmn_intron_F: 5’ ggatccCTGAAAATTTCCTTGCC 3’ XhoI_cmn_exon53/54_R: 5’ ctcgagccCTTGGTATTTATTTC 3’

To direct Cas9 and the donor plasmid towards the C-terminal intronic sequence of *cmn*, the following gRNA target site was used: 5’ ggccctgatttctcacgg 3’

The donor plasmid, *cmn* gRNA, and Cas9 protein were injected into zebrafish embryos at the single cells stage as previously described (Levic et al., 2021).

### Drug treatments

**Early DAPT treatment:** Transgenic embryos expressing *Bactin:pxna-GFP* and *entpd5a:pkRED* were incubated at 28°C for 7 hours. At this stage, embryos were dechorionated with a concentration of 1mg/mL of pronase dissolved in egg water and then treated with a 100mM concentration of DAPT for 1.5-3 hours (Özbudak & Lewis, 2008). DAPT treatments were washed out with three rinses of egg water and embryos were left to develop at 28°C until they were transferred to the aquaculture system at 5 dpf. Larvae were imaged starting at 5 dpf with either the AX10 Zoom V116 Zeiss microscope or SP8 Leica Confocal microscope.

### Microscopy

**Live imaging:** Whole-mount confocal imaging was performed on an SP8 inverted confocal microscope (Leica) equipped with a 25×/0.95 Fluotar VISIR water objective and Leica Application Suite software (Leica). Fish were mounted on to glass bottom dishes in a 0.6% mixture of low melt agarose and egg water. Additional imaging was done using a AX10 Zoom V116 Zeiss microscope equipped with a Plan Neofluar Z 1x objective and Zen software (all from Carl Zeiss).

**Laser-cut injuries and imaging:** Laser-cutting injuries were performed using a spinning disk confocal head (Perkin Elmer) attached to a Zeiss Axio Imager.M2m microscope equipped with a 40×/1.0 NA W Plan-Apochromat dipping lens objective and a Hamamatsu EM-CCD (C9100) camera (Kiehart et al., 2006). Micro-Manager software was utilized for time lapse image acquisition and lasercutting (Open Imaging, San Francisco, CA) (Edelstein et al., 2010). Laser cuts were performed using a Nd:YAG UV laser minilite II (Continuum, San Jose, CA) at a power of 1.3-2.3 µJ (Nova Ophir power meter) with a steering mirror for precise laser incisions on larvae mounted in a mixture of 0.8% low melt agarose and egg water.

### Image processing and analysis

**Image processing:** When necessary, images were minimally processed in ImageJ software (National Institutes of Health) for brightness and contrast. Digital stitching of confocal images was done in the Leica Application Suite (Leica) software. To generate surface projections, an open-source MATLAB toolbox ImSAnE (Image Surface Analysis Environment) customized for cylindrical shapes was utilized (Heemskerk & Streichan, 2015).

**Ordering index determination:** Images were pre-processed using a Gaussian blur operation followed by background subtraction in ImageJ. A binary mask of the notochord was then generated manually for thresholding.

First to avoid the continuous line of ventral expression prior to cluster formation in *entpd5a+* embryos, binary masks were slightly eroded. Then, we applied a fixed threshold to each imaging time point to extract the notochord segments clusters. For all experimental conditions, (wild-type, *fss^-/-^* or DAPT treated) the first two time points of the time course use a higher threshold of 25 to avoid expression from other notochord cells, since segment expression was dim. A threshold of 15 was then applied to all the other images (Figure S2B). Connected regions of binary pixels were identified as an *entpd5a*+ cluster, or a segment. For each cluster, the total number of pixels was used as a metric for segment area.

Cluster or segment areas were used to calculate an ordering index, which was a ranked list of sorted areas (Figure S2C). This sorted list then determined the progression of segment formation. We used several imaging time points taken over the same region to build a complete picture of the dynamics. For each subsequent time step, information about the order along with the center of mass locations for each cluster were compared to the previous time step.

For identifying new clusters not detected in the previous image, we used pairwise cluster-to-cluster distances along the AP axis, which we found remained fairly constant over time (Figure S2D). We compared pairwise distances over two subsequent timepoints - if new clusters filled in between two segments, the pairwise distances in the new image would not align perfectly with previously recorded distance (Figure S2E). To account for this change, the new cluster-to-cluster distances were summed until the results were similar. The number of new pairwise distances that needed to be added to match the previous pairwise distance identified the number of new clusters at that time point. Finally, any leftover pairwise distances indicated clusters formed to the right of the last known segment. Following our pairwise comparison, these new clusters were again ranked according to their area and added to the running list of clusters. Notably, we found that simply using cluster positions was ineffective as the tissue elongated over each time point causing the segments to shift.

Mature segments, especially in *fss* mutants, often merge, potentially affecting the calculation of ordering index by incorrectly assigning pairwise distances. To avoid this, we continuously identified regions (beginning from the anterior) that were already patterned and could not add any new segments as they had no more space to fill in. Once these regions were determined, segment areas and positions were no longer updated in the subsequent time points. This did not affect the index calculation as fixed clusters would already have been sorted and added to the running list. The final cluster in this region was then aligned using image registration with the next image for the calculation of pairwise distances and this process was repeated.

After the calculation of the ordering index, deviations from an expected sequential pattern were calculated (Figure 2D-F). Additionally, spacing between two ranked segments was also recorded from the ranked list of segments (Figure S5A, Figure 6D).

**RFP-Vinculin cluster analysis:** To extract Vinculin puncta, images were first processed through a small median filter. This effectively treated the puncta as noise, removing them. This processed image was then subtracted from the original image to remove all background, leaving only the puncta signal. Next, since each fish had slightly variable expression levels of RFP-Vinculin protein, we determined separate thresholds for each individual image quantitatively. This was done by plotting fluorescent intensity distributions for each image and converting them to a probability map of the cumulative intensity sum. The threshold for each fish was then chosen at the intensity value which corresponded to 0.997% of the sum. After thresholding, the images were then passed through ‘dbscan’ in MATLAB. This function performs a neighbor search and generates cluster cores based on a given radius and number of points within the specified region. This radius and number of points that defined a cluster of puncta was kept the same for each fish. The center of masses of the clusters were then extracted and compared to the segment locations at two consecutive time points of image acquisition.

**DN-Yap expression analysis:** Pre-processing steps of Gaussian blurring and background subtraction were applied as before for extracting segment clusters. A fixed threshold was then used to determine a segment. Areas per cluster across the AP axis were collected and grouped together to compare between wild-type and injected fish.

**Segment seeding analysis:** Surface projections were generated using the open-source MATLAB toolbox ImSAnE (Image Surface Analysis Environment). Images were then segmented using a fixed threshold after pre-processing with a Gaussian blur operation and background subtraction. For segment initiation events, only clusters below a cut-off area value were chosen to be seeds.

Each cluster’s center of mass was calculated along the Dorsal-Ventral-Dorsal Axis (*y_c_*) from surface projections. Using the total length of this surface projection axis (*L*), we recalculated the radial locations of the seeds around the notochord tube as:

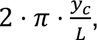 with *π* representing ventral regions and 0 and *2 ⋅ π* dorsal regions.

### Theoretical Modeling

Using as phenomenological order parameter Φ, which characterizes whether a cell of the notochord has been activated by Notch (Φ > 0) or not (Φ < 0), we describe the experimental system with a spatially anisotropic Brazovskiǐ effective free energy functional,

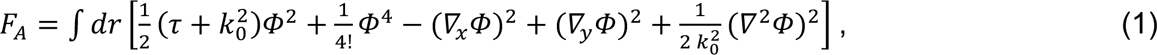

and its corresponding dynamics,

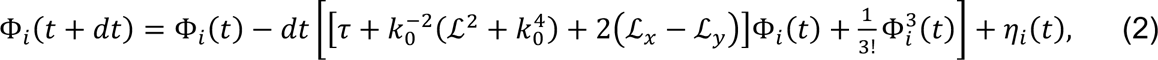

where *ℒ* is the Laplacian. Equation (2) is here discretized at the cell level, i.e., with mean cell width Δx = Δy = 1, for L_x_ = 192 and L_y_ = 16, under periodic boundary condition along the y-axis. These choices recapitulate the geometry of the notochord as a cylinder with L_y_*≪*L_x_, and results in the formation of 18-20 segments, as in experiments. The small noise strength chosen, 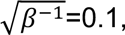 also recapitulates the sharp wall boundaries between *entpd5a* and *col9a2* domains. The presence of a crossover between nucleation and front propagation regimes as noise increases, however, is largely independent of β (not shown here). Under these conditions, a fixed time step of dt = is found to be sufficiently small to make the integration results invariant.

To evaluate how spatial disorder affects the patterning dynamics, we also consider

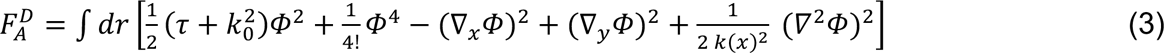

For σ = 0, *F*_A_ and 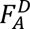 are identical, but generally σ > 0. Experimental estimates of σ are obtained by first extracting the mean cell-width (in μm) and the intensity profiles along the *x*-axis, *I(x)* vs *x* for each of the WT and *fss* fish using ImageJ (Figure 7C-F). The *x* axis is then rescaled by the average width of a cell (also obtained in μm) in the sample image, thus mapping experimental values on the model. Finally, the Fourier transform (FT) of the rescaled spatial profile is fitted to a normal distribution,

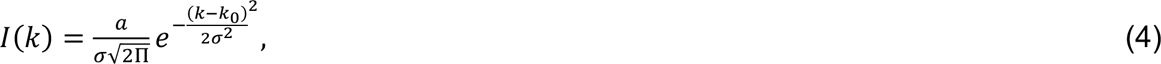

*k_0_* being the mean and σ the standard deviation of the distribution. A normalizing amplitude *a* is used to scale the distribution over the relevant range. Note indeed that the low *k* behavior (especially for *fss*, see Figure 7I-J) reflect the offset of the signal *I(x)* around their mean value, and hence do not meaningfully reflect the fluctuations in *k_0_*. Therefore, *I(k)* is only fitted over the range *k = (0.5, 1.1)*.

---

**Supplementary Figure 1:**
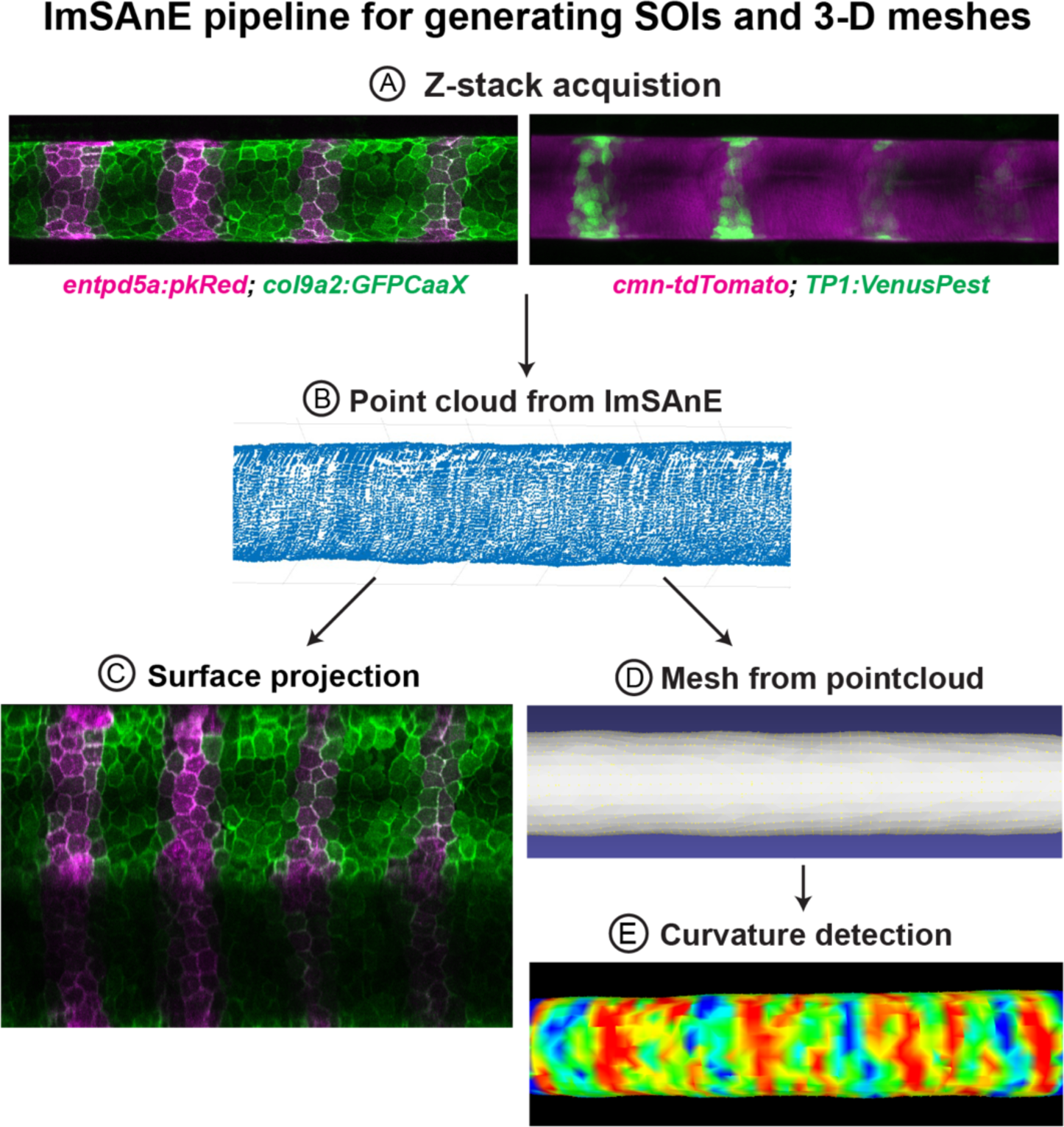
ImSAnE processing to generate surfaces of interest (SOIs) and 3-D meshes. A) Z-stack acquisitions capture the entire surface of the notochord. B) ImSAnE generates a point cloud from surface detection scripts and Ilastik training sets. C) Sample pullback of notochord sheath (same image as in A). D) Mesh generated in MeshLab, computed using exported point clouds. E) Computed Gaussian curvatures using the quadratic fitting method developed by MeshLab.

**Supplementary Figure 2:**
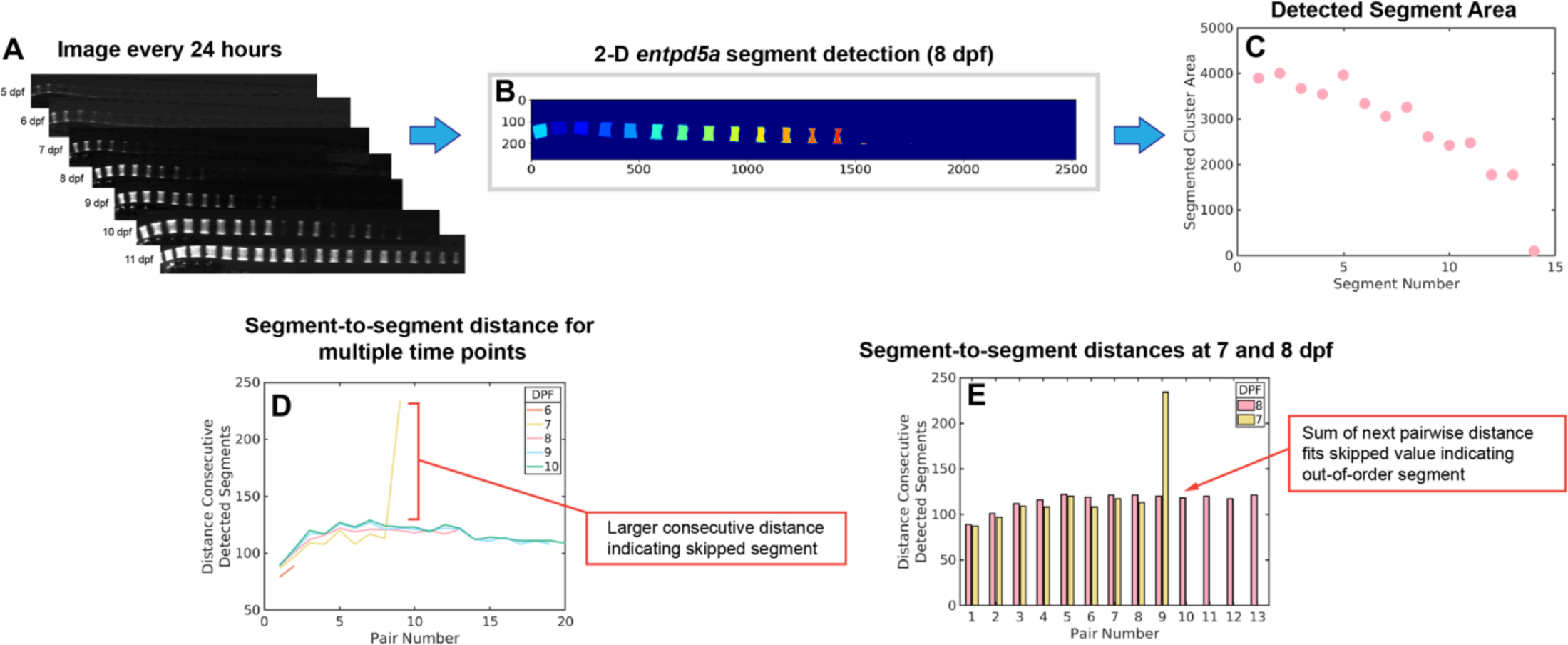
Segment detection for determining ordering index and segment-to-segment distances. A) Wide-field fluorescent images collected every 24 hours. B) 2-D segment detection. C) Plot of detected segment area. D) Segment-to-segment distances plotted for multiple time points. A large fluctuation in consecutive distances (orange bracket) indicates a skipped segment. E) Pairwise comparison of consecutive time points illustrates how consecutive segment distances are computed.

**Supplementary Figure 3:**
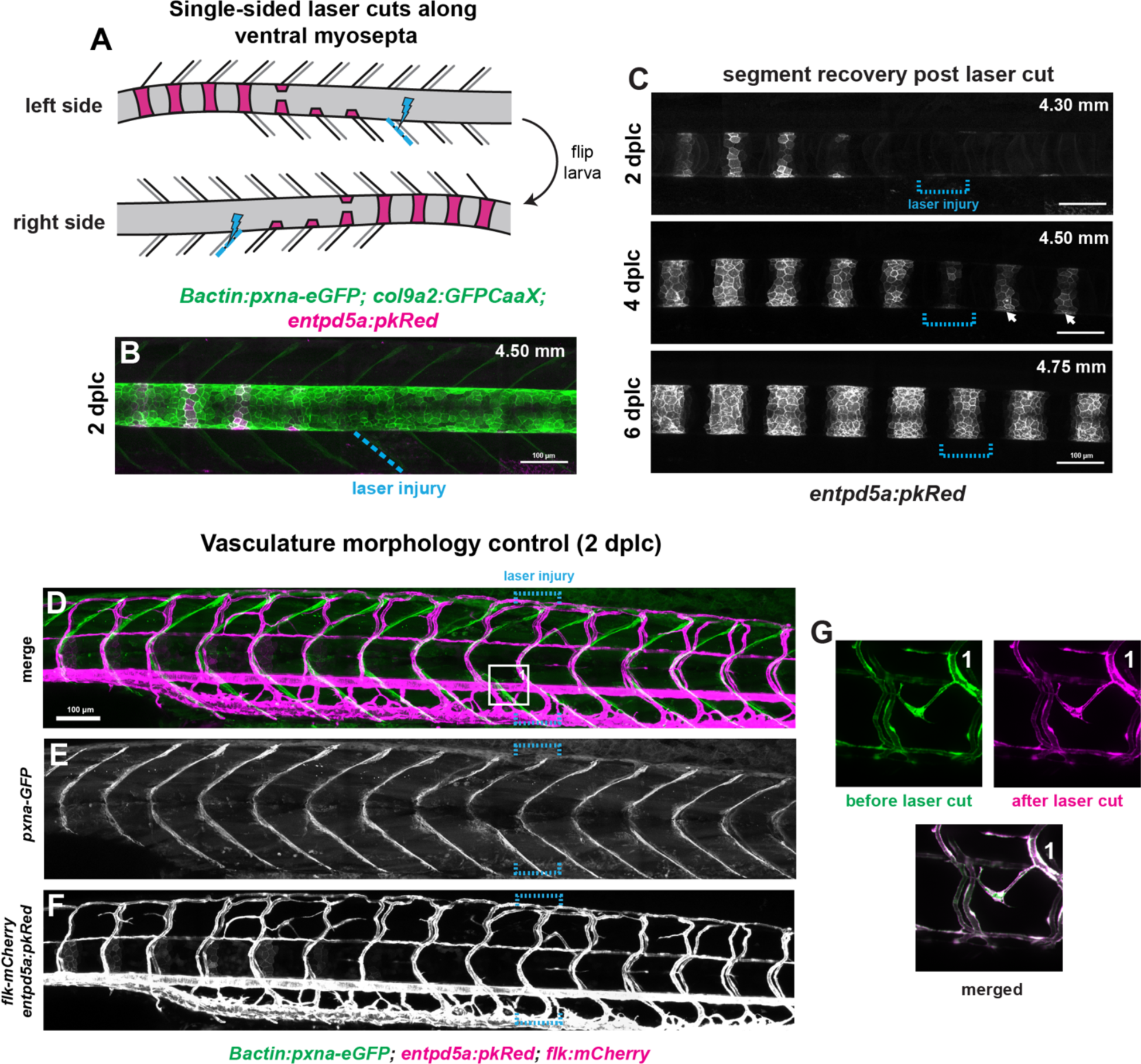
Laser cutting of ventral myosepta locally disrupts segment initiation. A) Schematic illustrating the bilateral laser cutting approach utilized to disrupt only ventral vertical myosepta (ventral-left, ventral-right). Injuries were always made in unsegmented regions in order to remove the segment initiation cue. B) Max intensity projection of a *Bactin:pxna-GFP*, *col9a2:GFPCaaX*, and *entpd5a:pkRed* larva injured 2 days prior. The dotted blue line indicates the location of the ventral. C) Time course of segment recovery following ventral myosepta cuts at 2, 4, and 6 dplc. The region of laser-injury is indicated by dotted blue brackets. At 2 dplc, the notochord segments adjacent to the injured region have not yet formed. At 4 dplc, notochord segments have started to form in the injured area, but the segment adjacent to the injured region is still delayed compared to more posterior segments (white arrows). By 6 dplc, the delayed segment has fully recovered and is nearly indistinguishable from segments in neighboring, unaffected regions. Scale bars = 100 µm. D) Merged confocal image of a larva expressing the vasculature reporter *flk:mCherry*, *entpd5a:pkRed*, and *Bactin:pxna-GFP* at 2 dplc. Bilateral laser-cuts were made along the myosepta denoted by blue brackets. E) Injured myosepta clearly show slackened morphology. F) Vasculature morphology in injured region is unaffected. G) Image panel shows max-intensity projections of the vasculature prior to performing the laser ablation on the adjacent myoseptum (green) and after injury (magenta). Overlay of the max intensity projections shows no immediate defects in vasculature morphology following injury.

**Supplementary Figure 4:**
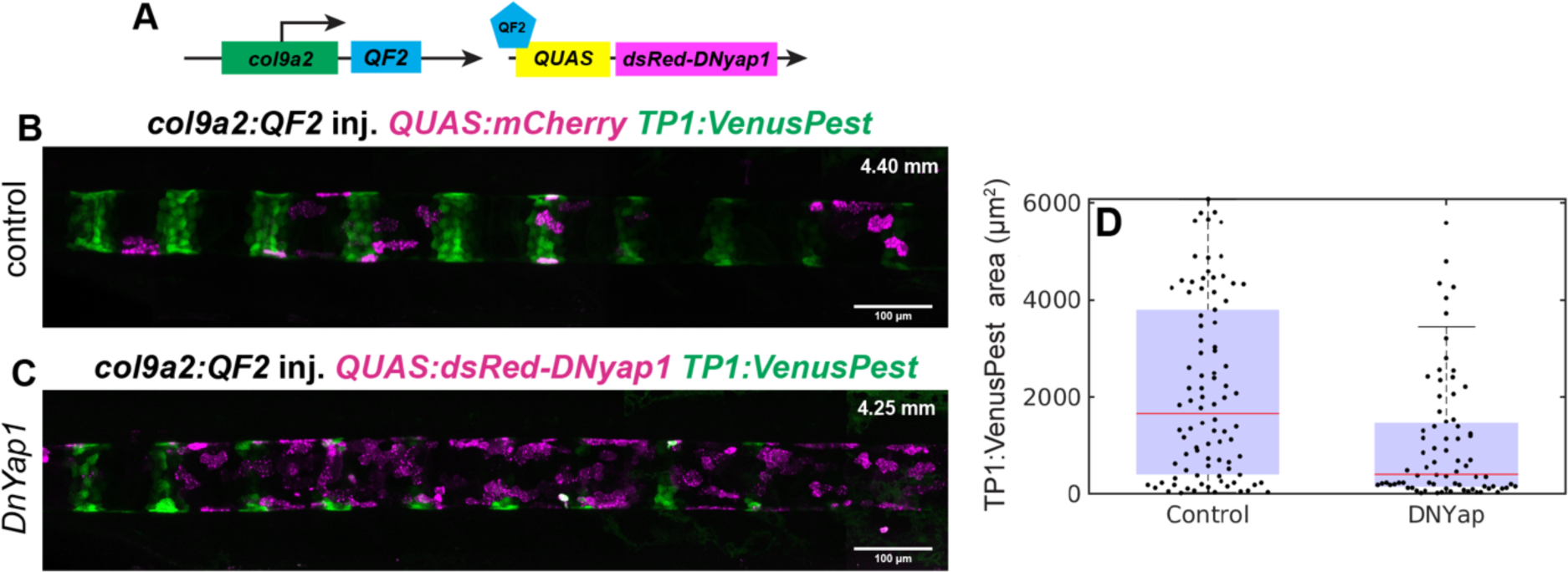
Inhibiting mechanosensitive transcription factor Yap halts notochord segmentation. A) Schematic of QF2/QUAS split system. We used a stable transgenic line to drive expression specifically in the notochord sheath. QUAS:dsRed-DN-yap or QUAS:mCherry control plasmid was microinjected at the 1-cell stage. B) Representative image of an 8 dpf larva expressing both *TP1:VenusPest* and *col9a2:QF2*, injected with *QUAS:mCherry* plasmid. *TP1:VenusPest* expression displays normal segmented patterning. C) Representative image of an 8 dpf larva expressing both *TP1:VenusPest* and *col9a2:QF2*, injected with *QUAS:dsRed-DNyap* plasmid. *TP1:VenusPest* expression is attenuated in regions of high DN-Yap expression. Scale bar = 100 µm. D) Plot depicting TP1 area (µm^2^) in either control (inj. *QUAS:mCherry*) or inj. DN-yap larvae (p value = 0.0016).

**Supplementary Figure 5:**
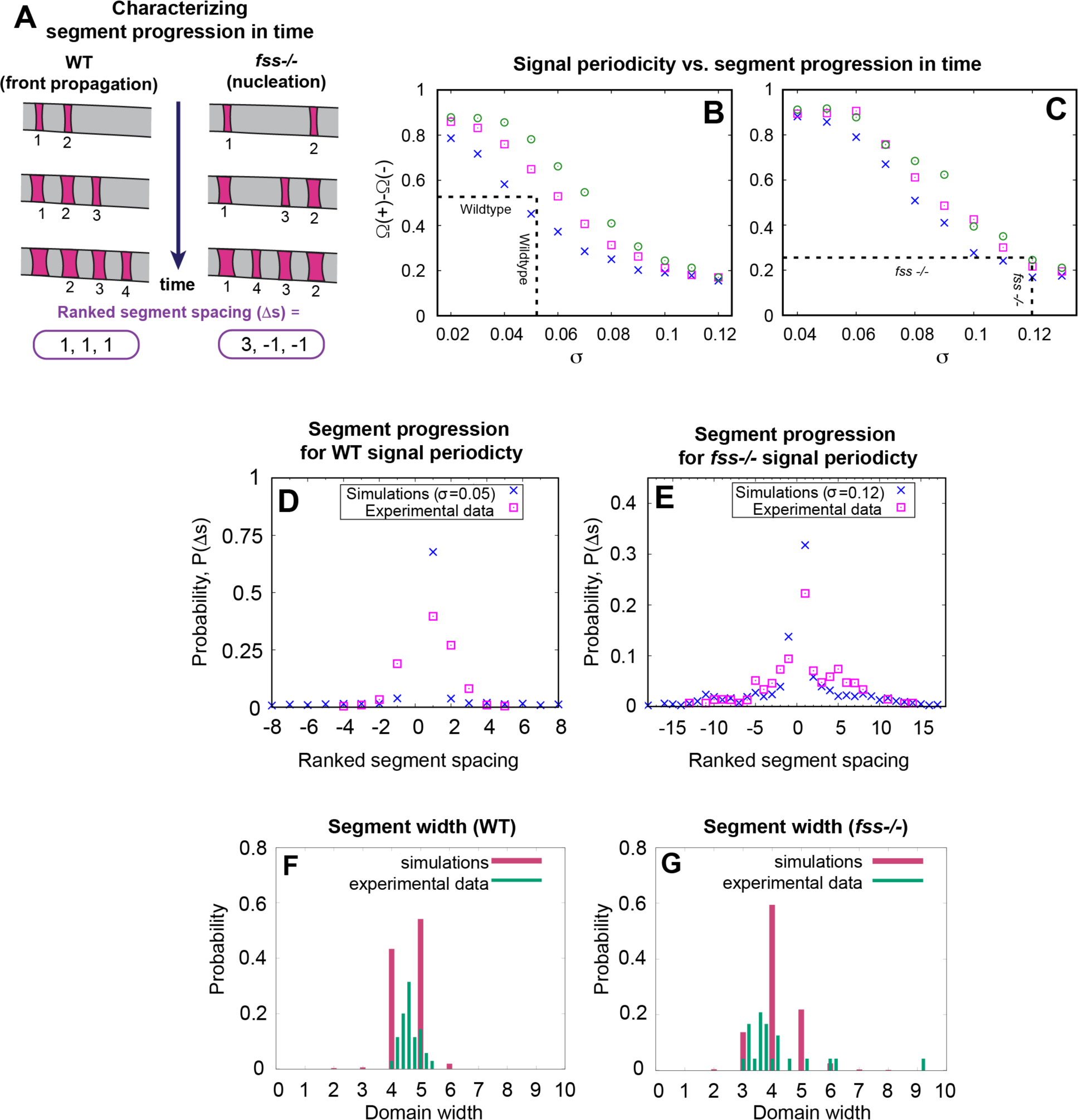
Comparing experimental and simulated segment ordering and width. A) Schematic illustration of ranked segment spacing (Δs) and their assigned values. Consecutively formed segments are assigned values Δs=1, whereas out-of-order segments receive higher or lower values. B) Plot estimating the value of *τ* in WT. Ω(+) denotes the sum of the probability for all positively ranked segments and Ω(-) denotes the sum of the probability for all the negatively ranked segments. Ω(+) and Ω(-) are shown for 3 different values of *τ* = −0.75 (blue crosses), −0.76 (magenta squares), −0.77 (green circles) at different values of σ. The difference, *Γ*=Ω(+)-Ω(-) obtained from simulations for different values of σ is then compared with that of experimental values of *Γ*_WT_ by using *σ*_WT_ (dashed lines) to obtain *τ*. This method gives *τ*_WT_ = −0.76 (see Supplementary Text). C) Plot estimating the value of *τ* for *fss* mutants by calculating from simulations and experiments using 3 different *τ* = −0.78 (blue crosses), −0.79 (magenta squares), −0.80 (green circles). *Γ* obtained from simulations for different values of σ is then compared with that of experimental values of *Γ*_fss_ by using *σ*_fss_ (dashed lines) to obtain *τ*. This method (see Supplementary Text) gives *τ*_fss_ = −0.80. D) Plot comparing the probability of a given ranked segment spacing, P(Δs) from the experimental data for WT and simulations at σ_WT_=0.05, *τ*_WT_ = −0.76. E) Plot comparing the probability for a given ranked segment spacing, P(Δs), from the experimental data for *fss*-/-and simulations at σ_fss_=0.12, *τ*_fss_ = −0.80. F) Using the same small σ value = 0.05 (see Table I, Supplementary Text) to model WT spatial periodicity, we find segment width clusters around 4-5 cells. For WT, we use k_0_ = 0.7 (see Table I, Supplementary Text). G) Using the larger σ value = 0.12, derived from the variation observed in *fss -/-* signal periodicity (see Table I, Supplementary Text), we observe a broader distribution of segment widths. For *fss -/-*, we use k_0_ = 0.75 (see Table I, Supplementary Text).

## Supplementary movie legends

**Supplementary Movie 1:** Simulation of the front propagation regime when σ = 0.05.

**Supplementary Movie 2:** Simulation depicting small loss of order when σ = 0.05.

**Supplementary Movie 3:** Simulation of non-sequential ordering (nucleation regime) when σ = 0.12.

