## Supplementary figures and images for "Axial segmentation by iterative mechanical signaling"

### Supplementary Movie 1

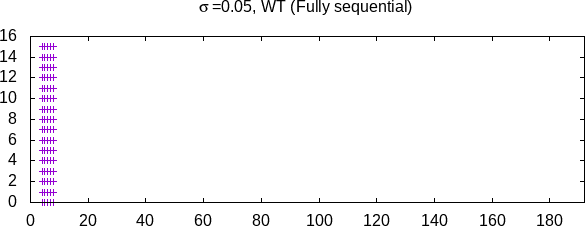

### Supplementary Movie 2

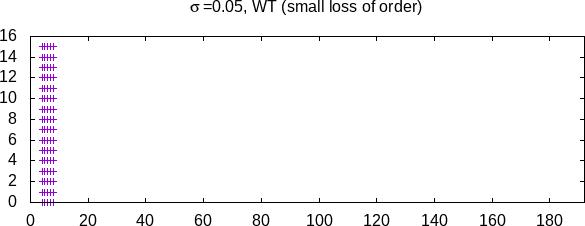

### Supplementary Movie 3

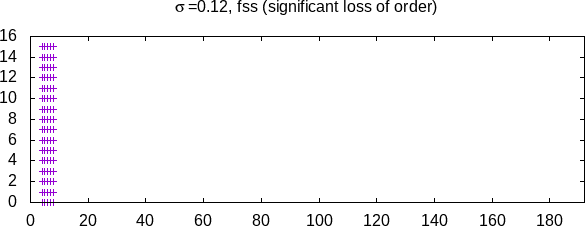
