## Supplementary Text for "Axial segmentation by iterative mechanical signaling"

We propose a coarse-grained model that recapitulates the striped domain geometry and their propagation during notochord segmentation. The equilibrium patterns are described by a frustrated effective free energy functional for a scalar order parameter Φ, which is positive for a cell with active notch signaling and negative otherwise, and for a given level of noise, $\tau$ . At low $\tau$, the free energy functional is minimized by the creation of interfaces between domains of a length scale, $\lambda = 2\pi/ k_0$. The dynamics obtained from an associated Langevin equation then results in domain formation crossing over from nucleating (non-sequential segment formation) to front propagating (sequential) as $\tau$ diminishes. Larger λ favors front propagation. When somite boundaries are globally disrupted, a situation akin to a strongly spatially fluctuating $k_0$, nucleation dominates. This theoretical model suggests that regularly spaced and properly formed somite boundaries instruct sequential and regular notochord segmentation in a process akin to phase ordering.

#### *I. A COARSE-GRAINED MODEL FOR NOTOCHORD PATTERNING*

In order to characterize the state of the notochord, we define a phenomenological order parameter Φ, which captures whether a cell has been activated (Φ > 0) by notch or not (Φ < 0). The "equilibrium" (steady-)state of the tissue is a spatially modulated striped phase with alternate domains expressing *entpd5a* and *col9a2* genes. In quite general terms, we describe this system by a modified Brazovskiĭ free energy functional (Brazovskiĭ SA, 1975; Geissler & Reichman, 2004; Gross et al., 2000) in two dimensions

$$F_A = \int dr \left[\frac{1}{2}(\tau + k_0^2)\Phi^2 + \frac{1}{4!}\Phi^4 - (\nabla_x\Phi)^2 + (\nabla_y\Phi)^2 + \frac{1}{2\,k_0^2}(\nabla^2\Phi)^2\right] \quad (1)$$

where the x-axis corresponds to the anterior-posterior axis of the notochord, and τ is a measure of fluctuations in the system. During the notochord segmentation, domains are favored and inhibited in directions perpendicular to the x- and y-axes, respectively. We thus accordingly construct an effective free energy, with $-(\nabla_x\Phi)^2$ favoring interfaces perpendicular to x-axis and $+(\nabla_y\Phi)^2$ inhibiting domain formation perpendicular to the y-axis. The anisotropy inherent of the tubular form of the notochord makes plausible such distinction. The domain periodicity, $\lambda = 2\pi/ k_0$, is stabilized by the $\nabla^4$ term. For $\tau < 0$, domain formation is possible, while only a disordered phase is possible at $\tau > 0$. In the thermodynamic limit, a phase transition at $\tau = 0$ separates the two regimes (Brazovskiĭ SA, 1975; Geissler & Reichman, 2004).

We also consider whether the relaxation properties of the coarse-grained model could reproduce the patterning dynamics of the notochord epithelium. To do so, one must first define the initial state. Recall that at very early stages only the *col9a2*+ (Φ < 0) are expressed in the notochord and as time progresses *entpd5a*+ domains (Φ > 0) grow as the expense of *col9a2*+ expressed cells. This process is initiated by the formation of a seed of *entpd5a*+ at the anterior end, and additional *entpd5a*+ domains form sequentially. We therefore take the initial seeded state of the notochord at time t = 0 to be

$$\begin{aligned}\Phi(x) &> 0 \; if \; x_1 < x < x_2, \\ &< 0 \; otherwise, \end{aligned} \quad (2)$$

where $x_1$, $x_2 \ll L_x$, and $L_x$ and $L_y$ are the dimensions of the notochord. It should be pointed out that the choice of the initial state is key to determining the relaxation dynamics. For example, for a homogeneous and well-mixed (paramagnetic) initial state–in which Φ(x) randomly take positive or negative values–phase ordering is spontaneous with domains forming almost instantaneously. This random activation, however, does not represent the state of the notochord at any stage. By contrast, a ferromagnetic initial state (seeded as in Eq. (2)) is more reflective of the biological situation. (As we will see below, this construction also reproduces the essence of the experimental observations.) Following a quench from this out-of-equilibrium state, the time evolution of the order parameter is then captured by the Langevin equation,

$$\frac{\partial \Phi}{\partial t} = -\frac{\delta F_A}{\delta \Phi(r)} + \eta(t), \qquad (3)$$

where η(t) is the source of Gaussian white noise with zero mean and variance

$$\langle \eta(r_i, t_m)\eta(r_j, t_n)\rangle = 2\beta^{-1}(dt/\Delta x^2)\delta_{ij}\delta_{mn}. \qquad (4)$$

In the numerical implementation, we approximate the Langevin equation using the discrete equation,

$$\Phi_i(t+dt) = \Phi_i(t) - dt\left[\left[\tau + k_0^{-2}(\mathcal{L}^2 + k_0^4) + 2(\mathcal{L}_x - \mathcal{L}_y)\right]\Phi_i(t) + \frac{1}{3!}\Phi_i^3(t)\right] + \eta_i(t), \qquad (5)$$

where $\mathcal{L}$ is the discrete Laplacian.

### II. RELAXATION REGIMES: FRONT PROPAGATION AND NUCLEATION

Simulation results give that for k0 fixed and uniform over the whole system, the phase ordering kinetics starting from the unstable initial state given by Eq. (6) can occur via one of two mechanisms: a) front propagation giving rise to sequential domain formation, and b) nucleation giving rise to domains forming randomly over the system length. To calculate the velocity, V of the growing/moving front, we measure the time $t_f$ taken for two new layers to appear adjacent to the anterior-most seed. V is then given by V = $\langle 2\lambda/t_f \rangle$ where $\langle \cdots \rangle$ denotes averaging over realizations. The nucleation process, by contrast, is initiated by a seed forming randomly at any position within the system. If the seed is greater than a critical size, it then grows to form an entire layer. In order to estimate the nucleation rate, we calculate the time $t_N$ taken to form a layer (Φ(x) > 0 at all cells within the layer) of typical width λ/2 starting from an initial state where Φ < 0 for all the lattice sites within the system. The nucleation rate is then given by $R_N = \langle 1/t_N \rangle$. We expect the nucleation to dominate at high fluctuations (low $-\tau$) and front propagation to take over as the fluctuations decrease (high $-\tau$ ). More specifically, the crossover corresponds to the front coverage rate, V /$L_x$ exceeding that of the nucleation rate, $R_N$ .

### III. SPATIAL DISORDER: MODELING INTERACTION WITH SOMITES AND ESTIMATING $k_0$ AND σ FROM THE EXPERIMENTAL DATA (ALSO SEE FIGURE 7 FROM MAIN TEXT)

Experimental results indicate that notochord segment initiation is strongly influenced by somite boundaries. When somites are disrupted, such as in *fss* mutants, notochord patterning becomes markedly imprecise. The front propagation mechanism is then not robust. Instead, pattern formation proceeds non-sequentially, in a way reminiscent of nucleation. In order to recapitulate these experimental observations, we consider models with spatially disordered frustration, *k*. The value of k at a specific site is then drawn from a Gaussian distribution *f(k)* with mean $k_0$ and standard deviation σ. The free energy in this scenario is then modified as

$$F_A^D = \int dr \left[\frac{1}{2}(\tau + k_0^2)\Phi^2 + \frac{1}{4!}\Phi^4 - (\nabla_x \Phi)^2 + (\nabla_y \Phi)^2 + \frac{1}{2\,k(x)^2}\,(\nabla^2 \Phi)^2\right] \quad (6)$$

Note that even though $k_0^2$ appears in the co-efficient of $\Phi^2$, we still keep it as a constant in $F_A^D$, because this coefficient measures the distance from the critical point. All the spatial dependence is therefore incorporated within the co-efficient of the $\nabla^4$ term.

Estimates for *f(k),* $k_0$ and σ are obtained from experiments. Taking the position x along the AP-axis, we first obtain the intensity $I_0(x)$ vs *x* for the segment positions as well as the Paxillin localization for each sample. The Fourier transform (FT) of the intensity

$$I_0(k) = \int dx\, I_0(x) e^{ikx} \quad (7)$$

for wavevector *k* is obtained and averaged over $N_s$ samples $I(k) = \langle I_0(k)\rangle_{N_s}$, and then fitted to a normal form,

$$I(k) = \frac{a}{\sigma\sqrt{2\Pi}} e^{-\frac{(k-k_0)^2}{2\sigma^2}}, \quad (8)$$

where *a* is an overall scale factor, and $k_0$ and σ are mean and standard deviation of the distribution, respectively. Experimental values are reported in Table I. Here, we have used $N_s = 6$ while measuring *I(k)* corresponding to the *entpd5a*+ segments and paxillin for both WT and *fss* (Figure 7H-K). For various σ, the nucleation rates and the velocity of the front are measured in the same way as described in Section II

### *IV. CHARACTERIZING THE TEMPORAL ORDER OF SEGMENTS AND ESTIMATING THE PARAMETER $\tau$ (ALSO SEE FIGURE 5 SUPPLEMENTARY)*

In order to characterize the temporal order, we define $s_1, s_2 \cdots s_N$ as the positions of the first, second, . . . , and N-th segments to appear over time. (The sequence $s_1, s_2 \cdots s_N$ is in time, not in space.) Note that for a pattern that forms purely sequentially, one gets $s_1 < s_2 < \cdots s_{N-1} < s_N$ . We also define the ranked segment spacing, Δs, which is a measure of the number of segments occupied between any two successively (in time) formed segments (see Supplementary Figure 5A) . Here, we compute Δs from the experimental data (n=10 for WT and n=7 for *fss*) and also by simulating our model (for about 200 realizations). Therefore, for a fully sequential segment, Δs = $s_2 - s_1 = s_3 - s_2 = \cdots = 1$ for all the N −1 pairs. This property would no longer be preserved for segments formed out of temporal sequence. When Δs > 0, the next segment is formed towards the posterior side of a given segment, whereas when Δs < 0, the successive segment is located on the anterior side of the current segment. For brevity, we name as "forward" segments those with Δs > 0 and "backward" segments those with Δs < 0. The probability distribution, P(Δs) is then used as a good proxy for the temporal order. Now, in order to make an estimate of the parameter $\tau$, we measure, for various values of σ, $\Omega(+) = \sum_{\Delta s>0} P(\Delta s)$ (Supplementary Figure 5B-C), which is the sum of the probability of all the forward segments. Similarly, we compute $\Omega(-) = \sum_{\Delta s<0} P(\Delta s)$, which is the sum of the probability of the backward segments. The difference, $\Gamma = \Omega(+) - \Omega(-)$ is then computed from the numerical simulations of the model and then compared it with the experimentally obtained values of $\Gamma_{expt}$ for the WT and *fss*. Using σ = 0.05 as the reference value for WT and σ = 0.12 for *fss*, we estimate $\tau$ at which matches that from the simulations. This exercise gives us $\tau_{WT} = -0.76$ and $\tau_{fss} = -0.80$. We then use $\tau_{WT}$ , $\sigma_{WT}$ for the WT case and $\tau_{fss}$, $\sigma_{fss}$ for the fss case as reference values in our simulations to compare with the distributions P(Δs) obtained from the data of the sample images. As seen in Supplementary Figure 5D-E, the simulations results agree fairly well with the experimental data.

| | a | $k_0$ | σ |
|---|---|---|---|
| WT (*entpd5a*) | 0.012(3) | 0.70(1) | 0.05(1) |
| WT (*pxna*) | 0.012(3) | 0.70(1) | 0.07(2) |
| *fss* (*entpd5a*) | 0.012(3) | 0.75(3) | 0.12(4) |
| *fss* (*pxna*) | 0.009(3) | 0.87(7) | 0.27(7) |

**TABLE I.** Parameters a, $k_0$ and σ as determined from the Fourier transform analysis of WT and *fss* images within 95% confidence interval
